# CENP-A overexpression drives distinct cell fates depending on p53 status

**DOI:** 10.1101/2020.07.21.213496

**Authors:** Daniel Jeffery, Katrina Podsypanina, Alberto Gatto, Rebeca Ponce Landete, Lorraine Bonneville, Marie Dumont, Daniele Fachinetti, Geneviève Almouzni

## Abstract

Tumour evolution is driven by both genetic and epigenetic changes. CENP-A, the centromeric histone H3 variant, is an epigenetic mark that directly perturbs genetic stability and chromatin when overexpressed. Although CENP-A overexpression is a common feature of many cancers, how this impacts cell fate and response to therapy remains unclear. Here, we established a tunable system of inducible and reversible CENP-A overexpression combined with a switch in p53 status in human cell lines. Through clonogenic survival assays and single-cell RNA-sequencing over time, we uncover the tumour suppressor p53 as a key determinant of how CENP-A impacts cell state, cell identity and therapeutic response. If p53 is functional, CENP-A overexpression promotes senescence and radiosensitivity. But, when we inactivate p53, CENP-A overexpression instead promotes epithelial-mesenchymal transition, an essential precursor for tumour cell invasion and metastasis. Thus, CENP-A overexpression drives distinct cell fates depending on p53 status, with important implications for tumour evolution.

## Introduction

Tumour evolution is driven by both genetic and non-genetic changes from tumourigenesis to therapeutic resistance (Easwaran et al., 2014; Shen and Laird, 2013). The fate of a given cell depends on how these changes impact both cell state (e.g. proliferation or cell death) and cell identity (e.g. stemness or differentiation). On the genetic side, tumour evolution is strongly linked to chromosomal instability (CIN)(Gerstung et al., 2020; Tijhuis et al., 2019; Ye et al., 2018). This sub-category of genome instability is characterized by an increased rate of mutagenesis through losses or gains of chromosomes, partial chromosomes or chromosomal rearrangements. While extreme CIN is generally deleterious to cells, low or intermediate CIN increases the heterogeneity of the cell population (Potapova et al., 2013). These changes can promote the selection of advantageous clones through Darwinian evolution (Gerlinger and Swanton, 2010) a hypothesis put forward more than a century ago in cancer biology (Boveri, 2008). On the non-genetic side, environmental and metabolic insults can induce changes to epigenetic landscapes that perturb genome regulation (Flavahan et al., 2017). This can further contribute to tumour heterogeneity (Guo et al., 2019) enabling cell plasticity (Wainwright and Scaffidi, 2017), acquisition of stemness properties (Lytle et al., 2018) and the ability to counteract cell shutdown mechanisms (Flavahan et al., 2017). All of these aspects promote tumour development and reduce therapeutic response. But, in contrast to genetic changes, epigenetic changes are reversible (Feinberg and Fallin, 2015). This reversibility makes epigenetic marks and their regulatory factors interesting targets for cancer treatment, especially in combination with other anti- cancer therapies (Morel et al., 2019).

As an epigenetic mark with direct effects on CIN, the centromeric histone H3 variant, Centromere Protein A (CENP-A), is of particular interest (Sharma et al., 2019). In mammals, CENP-A is considered as a prime example of a bona fide transgenerational epigenetic mark (Das et al., 2017). Indeed, its deposition determines the location of the centromere (Barnhart et al., 2011; Fachinetti et al., 2013). Furthermore, it is transmitted across cell divisions and even organismal generations (Amor et al., 2004; Tyler-Smith et al., 1999). CENP-A is deposited (Dunleavy et al., 2009; Foltz et al., 2009) and maintained (Zasadzińska et al., 2018) at centric chromatin by its dedicated histone chaperone HJURP. From there, CENP-A acts as the foundation for kinetochore assembly during cell division, where it is essential for efficient chromosome segregation (reviewed in Fukagawa and Earnshaw, 2014; McKinley and Cheeseman, 2016; Musacchio and Desai, 2017). Therefore, the maintenance of genome integrity depends on the proper regulation of CENP-A (Sharma et al., 2019). This is demonstrated by CENP-A depletion (Maehara et al., 2010) and overexpression (Shrestha et al., 2017) experiments, which both promote mitotic defects and CIN in human cells.

Importantly, high CENP-A levels correlate strongly with increased tumour aggressiveness in patients, including increased tumour stage/size (Gu et al., 2014; Li et al., 2011; Ma et al., 2003; Qiu et al., 2013; Saha et al., 2020; Sun et al., 2016; Zhang et al., 2016), increased invasiveness/metastasis (Gu et al., 2014; Ma et al., 2003; McGovern et al., 2012; Saha et al., 2020; Sun et al., 2016; Zhang et al., 2016), increased rates of recurrence (McGovern et al., 2012; Sun et al., 2016; Zhang et al., 2016) and poor patient prognosis (Gu et al., 2014; McGovern et al., 2012; Qiu et al., 2013; Sun et al., 2016; Wu et al., 2012; Zhang et al., 2016). Examining the regulation of *CENPA* gene expression, we demonstrated that the tumour suppressor p53 (reviewed in Levine, 2020) negatively regulates CENP-A transcription through its downstream effector p21 (Filipescu et al., 2017). Thus, CENP-A levels should be held in check by active p53. But CENP-A overexpression occurs in many cancers (Saha et al., 2020; Sun et al., 2016), and although *TP53* mutation correlates with higher CENP-A levels in patient tumours (Filipescu et al., 2017), many tumours overexpress CENP-A despite having wild type *TP53* (cBioPortal: Cerami et al., 2012; Gao et al., 2013), indicating that p53 regulation is not the sole mechanism that controls CENP-A expression. Notably, correlation of high CENP-A levels with therapeutic response to DNA damaging agents is a matter of debate, with studies arguing for reduced or improved response (Zhang et al., 2016). However, p53 status was not assessed in these studies. This is even more important, given that the p53 pathway responds differentially to distinct cell-cycle defects (McKinley and Cheeseman, 2017). Thus, understanding how CENP-A overexpression impacts cell fate in different p53 contexts is an important question that could shed light on cell response to cancer treatment.

To address this issue, we set up a system to turn on and off the overexpression of CENP-A in various p53 contexts and switch p53 status in selected cell lines. Using this tunable system, we demonstrate that CENP-A overexpression alters cell fate in a manner dependent on p53 status. When p53 is functional, CENP-A overexpression alters cell state, promoting cell cycle arrest, senescence and radiosensitivity. But when we inactivate p53 the cells evade arrest. Instead, CENP-A overexpression drives a change in cell identity, promoting epithelial-mesenchymal transition (EMT). Thus, our findings reveal an unanticipated function of CENP-A overexpression to reprogram cell fate in distinct ways that depend on p53 status, with important implications for the role of CENP-A in tumour evolution.

## Results

### Inducible and reversible CENP-A overexpression in cells with varied p53 status

In order to explore how CENP-A expression levels and p53 status impact cell fate and cell response to anti-cancer treatment, we first established a system where we could specifically alter CENP-A expression independent of its regulation by p53. For this, we added a doxycycline-inducible CENP-A overexpression construct (*TetOn-CENPA-FLAG-HA*) by lentiviral transduction into several cell lines with varied p53 status (Figure 1A, see also Methods). Importantly, we did not use antibiotic selection to obtain CENP-A overexpression. This enables us to investigate the unbiased effects of CENP-A overexpression over time, without provoking cell adaptations or secondary mutations that could arise from selective pressure. The TetOn system allows induction of CENP-A overexpression within 24h of adding doxycycline (Figure 1B-D). This acute induced overexpression of CENP-A increases HJURP protein levels (Figure 1C), increases levels of CENP-A at centromeres and also causes mislocalization of CENP-A to the chromosome arms (Figure 1D), regardless of p53 status. These effects are similar to those observed with constitutive CENP-A overexpression in HeLa cells (Lacoste et al., 2014). Here, however we could reverse both CENP-A overexpression and mislocalization simply by washing out doxycycline (Figure 1C-D), consistent with the observation of rapid removal of ectopic CENP-A from chromosome arms with DNA replication (Nechemia-Arbely et al., 2019). Thus, we had at hand an efficient and reliable system to test therapeutic sensitivity in several cell lines with varied p53 status under conditions of tunable CENP-A overexpression.

**Figure 1.**
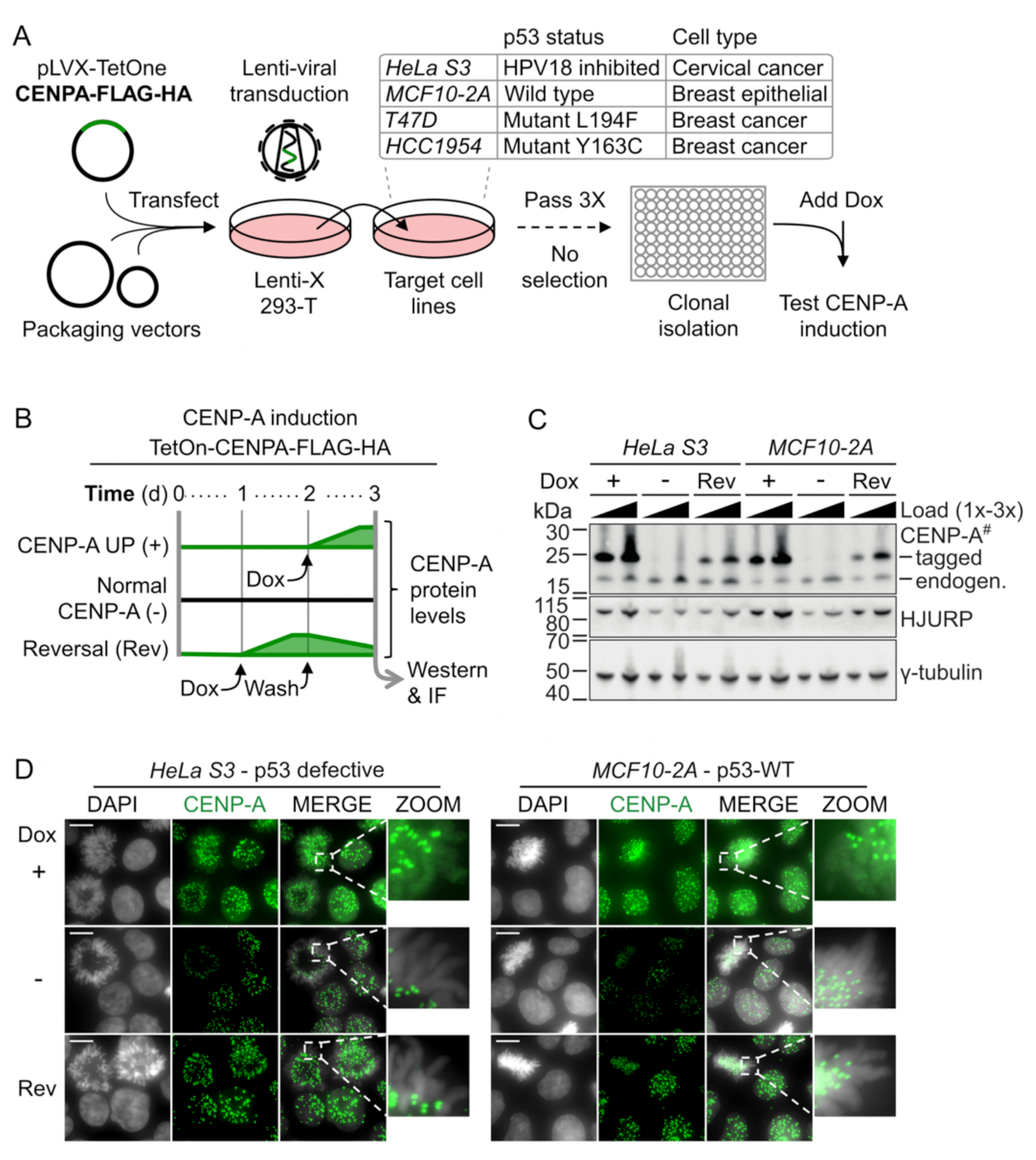
Inducible and reversible CENP-A overexpression in cells with varied p53 status. A) Scheme for the generation of doxycycline (Dox) inducible CENP-A overexpression cell lines. *TetOn-CENPA-FLAG-HA* construct was randomly integrated into several indicated target cell lines by lentiviral transfection without selection. After clonal isolation, we tested cells for clear homogenous CENP-A and HA increase by IF in approximately all cells 24h after the addition of 80ng/ml of Dox. Cells were also tested to ensure no detectable background HA signal by Western or IF, when no Dox was added. B) Scheme representing relative CENP-A protein levels over time for *TetOn-CENPA-FLAG-HA* cell lines. Dox = 10ng/ml. C) Western blot total cell extracts (TCEs) pertaining to scheme in B. Primary antibodies indicated on the right. Exogenous CENP-A (tagged) can be distinguished from endogenous CENP-A (endogen.) by its increased molecular weight. # = high sensitivity ECL. y-tubulin used as loading control. D) Immunofluorescence of paraformaldehyde-fixed cells using anti-CENP-A antibody (green) and DAPI staining (grey). Conditions in parallel with C. Showing max projection images from a Z-series with zoom on a mitotic cell (DAPI/CENP-A merge) to highlight chromosome arms. Scale bars = 10µm.

### Induced CENP-A overexpression causes reversible radiosensitivity in p53-WT cells

To test therapeutic sensitivity in our cell lines, we chose X-irradiation as a representative DNA damaging agent that is commonly used in cancer therapy. Thus, we tested cell survival after X-irradiation with or without induction of CENP-A overexpression by colony formation assays (CFAs), a surrogate for self-renewal capacity (Figure 2A). CENP-A overexpression in non-tumoural, WT p53 cells (MCF10-2A) led to a strong increase in sensitivity to X-irradiation. Meanwhile, CENP-A overexpression in all tumoural cell lines with defective p53 (HeLa S3, T47D, HCC1954 and DLD1) did not significantly affect radiosensitivity – a distinctly radio-tolerant status (Figure 2B, see also Figure S1A). In order to determine if the differences in sensitivity that we observed upon CENP-A overexpression were due to secondary mutations from mitotic defects, we tested the reversibility of this sensitivity by tuning down CENP-A expression to near-endogenous levels in the p53-WT cells (MCF10- 2A) and p53 defective cells (HeLa S3)(Figure 2C). Remarkably, restoration of CENP-A overexpression to endogenous levels reversed the sensitivity phenotype in MCF10-2A, while HeLa cells, used here as a control line, remained tolerant (Figure 2D). Therefore, the radiosensitivity observed in MCF10-2A cells requires the continued overexpression of CENP-A and it cannot be explained by genetic effects (i.e. secondary mutations) that occur after induced CENP-A overexpression. Thus, CENP-A overexpression leads to radiosensitivity in p53-WT but not p53-defective cells.

**Figure 2.**
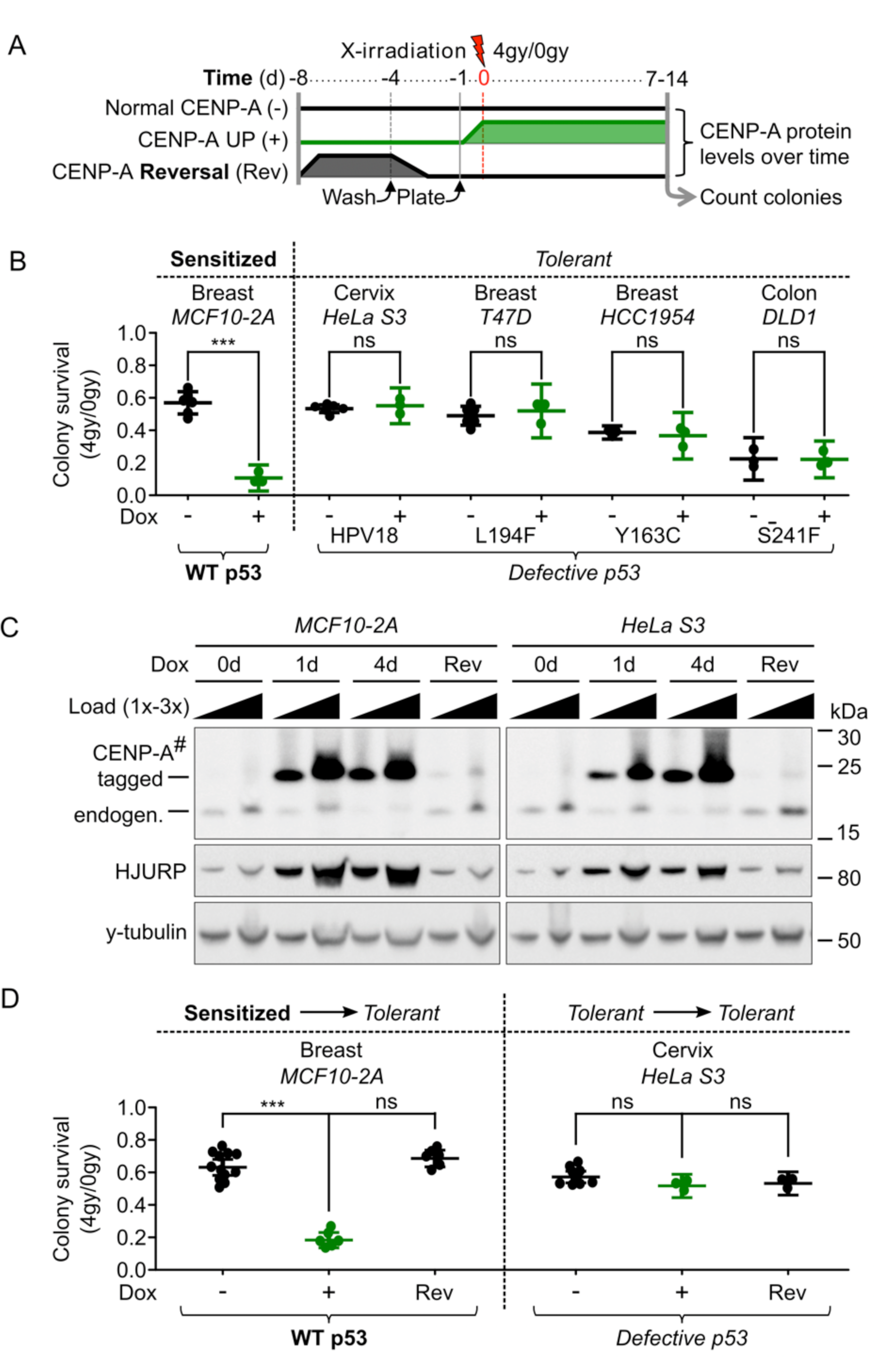
Induced CENP-A overexpression causes reversible radiosensitivity in p53 wild type cells. A) Scheme of colony formation assays (CFAs) with relative CENP-A protein levels over time for *TetOn-CENPA* cell lines. B) CFA results corresponding to the scheme in A for five different cell lines of varied tissue origins and p53 status, as indicated. Each dot represents a single biological replicate. Plots show mean ± 95% confidence interval. Statistical significance tested by two-tailed Welch’s t-test with Bonferroni cutoff at a p-value of 0.01 (α = 0.05). *** = p-value <0.0001. See also Figure S1A for survival ratios relative to untreated condition. C) Western blot corresponding to scheme in A: TCEs after 1 day (1d) or 4d with 10ng/ml Dox, no Dox control (0d), or 4d Dox followed by 4d without Dox (Rev). Primary antibodies indicated on the left. 1x load = ∼30000 cells. # = high sensitivity ECL exposure. y-tubulin used as loading control. D) CFA results corresponding to scheme in A for the indicated cell lines. Plots as in B. Statistical significance tested by two-tailed Welch’s t-test, compared to non-induced control, with Bonferroni cutoff at a p-value of 0.0125 (α = 0.05). *** = p-value <0.0001.

### Sensitivity to X-irradiation upon CENP-A overexpression is dependent on functional p53

To assess if p53 plays a causal role in the radiosensitivity associated with CENP-A overexpression, we took advantage of the possibility to switch the p53 status in two of our inducible CENP-A overexpression cell lines: the sensitized MCF10-2A cells and the tolerant HeLa cells. We could thus carry out CFAs after CENP-A overexpression and X-irradiation in isogenic cell lines, but with altered p53 status (Figure 3A). For the change from p53-WT to p53-defective, we stably transduced the clonal MCF10-2A *TetON-CENPA-FLAG-HA* cells with either an empty vector control or a constitutively expressed dominant negative (DN) *TP53* construct, in order to mimic p53 loss and p53 loss of function (LOF) mutations (p53DD construct, (Hahn et al., 2002)). The p53-DN peptide stabilizes the WT p53 protein, causing a clear increase in p53 protein level, but suppresses its activation of downstream targets (e.g. p21). Indeed, the indirect activator of p53 (Nutlin-3) caused clear increases in p53 and p21 in p53-WT cells, but not those expressing the p53-DN peptide (Figure 3B). For the change from p53-defective to p53-WT in the CENP-A inducible HeLa cells, we transiently activated WT p53 by incubation at 42°C for 1h immediately prior to X-irradiation. This treatment is known to temporarily release p53 repression from the HPV18 E6 protein, permitting a functional p53 response (Oei et al., 2015). Indeed, we confirmed that after hyperthermia treatment, p53 levels increased (Figure 3C), followed shortly after by increased p21 (Figure 3D). By switching p53 status, in each case we also altered the radiosensitivity of the cells. The p53-DN peptide in MCF10-2A cells strongly counteracted the radiosensitivity phenotype associated with CENP-A overexpression, while the short hyperthermia treatment in the HeLa cells was sufficient to mildly, but significantly, increase sensitivity to X-irradiation only after CENP-A overexpression (Figure 3E). We further confirmed the radiosensitivity phenotype using a range of doxycycline concentrations and a range of X-irradiation doses in CFAs for the p53-WT and p53-DN MCF10-2A cells (Figure S1B-C). Taken together, the results demonstrate that induced CENP-A overexpression increases radiosensitivity in a p53- dependent manner.

**Figure 3.**
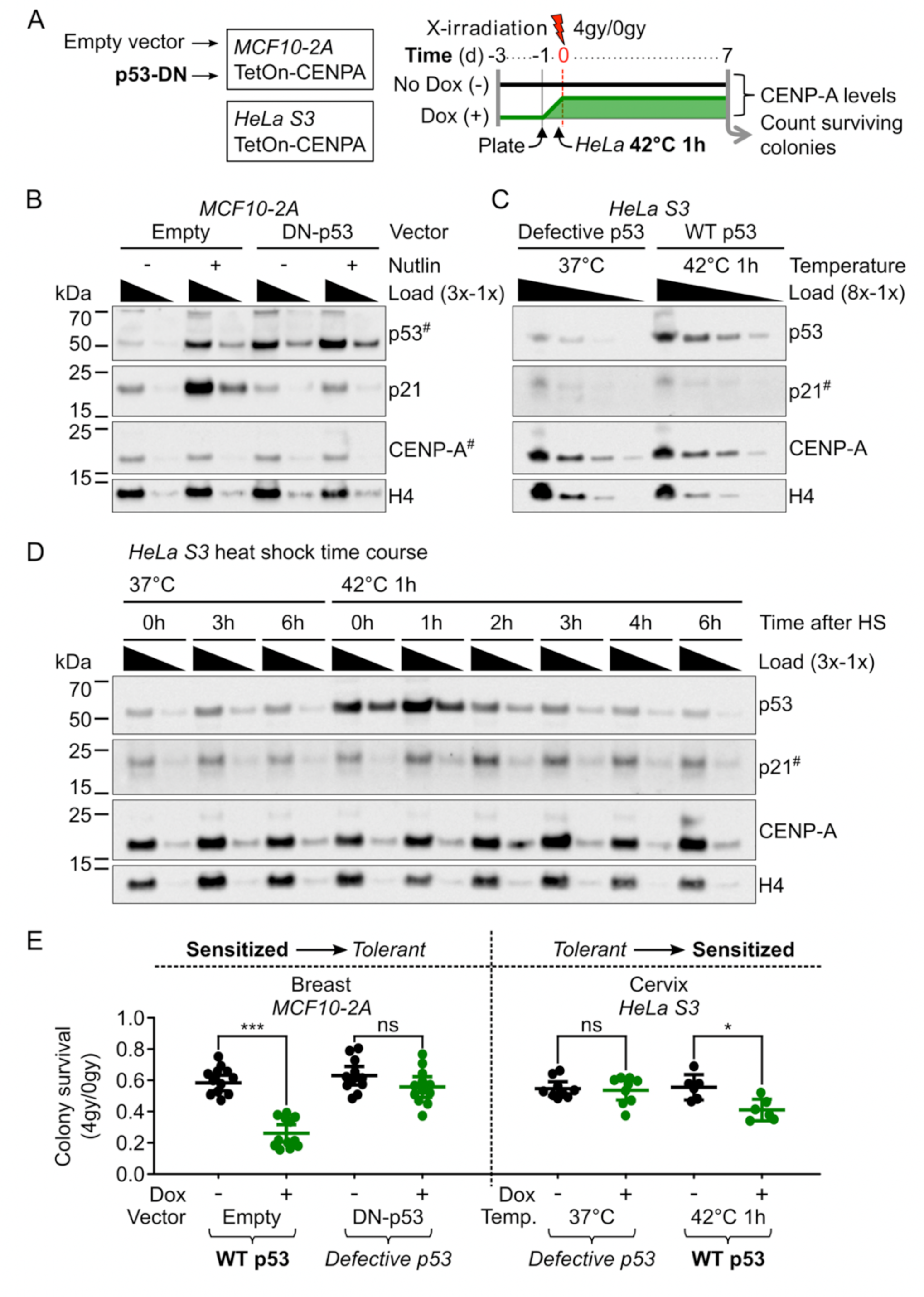
Sensitivity to X-irradiation upon CENP-A overexpression is dependent on functional p53. A) Scheme of CFAs and change of p53 status with relative CENP-A protein levels over time for *TetOn-CENPA-FLAG-HA* cell lines. B) Change of p53 status in MCF10-2A TetOn-CENPA-FLAG-HA cell. Western blot of TCEs after 24h of 10µM Nutlin-3 treatment or DMSO control, as indicated. Dominant negative p53 peptide stabilizes p53 and suppresses its activation of p21. 1x load corresponds to total protein extract from ∼30000 cells. Primary antibodies are indicated on the right. ^#^ = high sensitivity ECL exposure. Representative Western blot from three independent experiments is shown. H4 used as loading control. C) Increase of p53 levels in HeLa cells by heat shock. Western blot of TCEs after 1h at 42°C or 37°C control. Loaded at 8x, 4x, 2x, 1x concentrations, where 1x load = ∼12500 cells. Primary antibodies are indicated on the right. ^#^ = high sensitivity ECL exposure. H4 used as loading control. Representative Western blot from three independent experiments is shown. D) Temporary change of p53 in HeLa cells by 1h heat shock (HS). Western blot of HeLa cell TCEs at indicated time points following 1h at 42°C or 37°C control. Temporary increase of p53 and subsequently p21. 1x load = ∼33000 cells. Primary antibodies are indicated on the right. ^#^ = high sensitivity ECL exposure. H4 used as loading control. E) CFAs pertaining to scheme in A. Each dot represents a single biological replicate. Plots show mean ± 95% confidence interval. Statistical significance tested by two-tailed Welch’s t-test with Bonferroni cutoff at a p-value of 0.0125 (α = 0.05). *** = p-value <0.0001, * = p-value <0.01. See also Figure S1B-C for CFAs following range of Dox and X-irradiation doses.

### Induced CENP-A overexpression promotes radiosensitivity by impairing cell cycle progression

Given that the p53-dependent radiosensitivity was due to non-genetic (reversible) effects of CENP-A overexpression, we decided to assess changes at the transcriptional level. We first assessed global transcription by RNA-seq, analyzing the effects of acute CENP-A overexpression (at two different concentrations of Dox), switch of p53 status, and X-irradiation treatment in the MCF10-2A cells (Figure 4A). X-irradiation and inactivation of p53 affected the transcription of several genes (Figure 4B). Notably, CENP-A overexpression induced much more extensive changes, with more than 8000 genes affected. According to Gene Set Enrichment Analysis (GSEA), CENP-A overexpression mainly led to the repression of genes involved in cell cycle progression, DNA repair and RNA metabolism (Figure 4C, details in Table S1). To more specifically identify the key cellular pathways involved in radiosensitivity and its reversal by inactivation of p53, we applied hierarchical clustering to identify co-regulated subsets of differentially expressed genes (DEGs). In particular, we wanted to identify genes for which p53-DN could counteract the effects of CENP-A overexpression. Out of ten clusters (Figure 4D, see also Figure S2C), there was only one that corresponded to these criteria (Cluster 8). This subset of genes is strongly downregulated by CENP-A overexpression and upregulated by p53-DN (Figure 4E).

**Figure 4.**
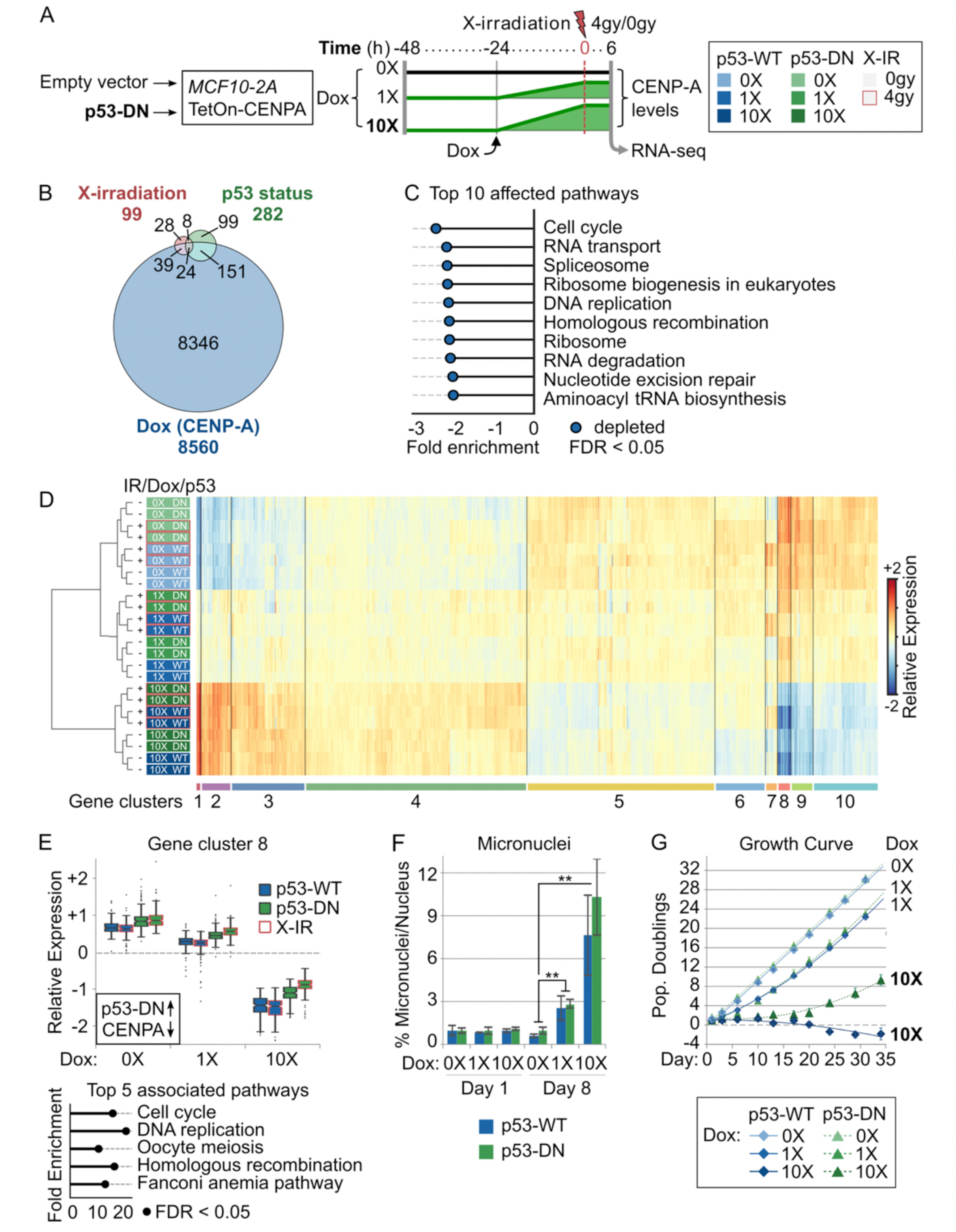
Induced CENP-A overexpression promotes radiosensitivity by impairing cell cycle progression. A) Scheme delineating conditions tested by RNA-sequencing showing relative CENP-A protein levels over time and corresponding legend (pertains to D and E). All conditions tested in duplicate. See Figure S1D for CENP-A protein levels corresponding to 0X, 1X (10ng/ml) and 10X (100ng/ml) Dox. The transcriptional impact of Dox treatment alone was tested in the non-inducible MCF10-2A parental control (See Figure S2A-B). B) Proportional Venn diagram summarizing the number of differentially expressed genes (DEGs) upon X-irradiation (Red, 0gy vs 4gy); change of p53 status (Green, p53-WT vs p53-DN); and CENP-A overexpression (Blue, 0X Dox vs 1X Dox vs 10X Dox). Number of DEGs in each category and overlapping categories is indicated. C) Gene Set Enrichment Analysis showing the top ten KEGG pathways affected by CENP-A overexpression (WebGestaltR v0.4.2) and normalized enrichment score. All ten pathways are significantly enriched in downregulated genes (i.e. depleted, at FDR < 0.05). See Table S1 (GSEA) for all significantly depleted/enriched processes. D) Heat map showing hierarchical clustering of samples (rows, coloured according to legend in A) based on the relative expression of all DEGs (columns), then classified into 10 main gene clusters (bottom). Expression relative to average of all genes (mean-centered counts, log_2_-transformed and TMM-normalized). E) Top: Box plots showing the distribution of expression levels for DEGs in Cluster 8 according to experimental condition. See Figure S2C for all gene clusters. Black box: for genes in this cluster, p53-DN increases expression (up arrow) and CENP-A overexpression decreases expression (down arrow). Bottom: Top five enriched KEGG pathways (WebGestaltR v0.4.2) for genes within Cluster 8 based on over-representation analysis (ORA). All five terms are significantly over-represented (FDR < 0.05). See Table S1 (ORA) for top ten pathways for all clusters. F) Accumulation of micronuclei pertaining to Day 1 and Day 8 of MCF10-2A *TetOn-CENPA- FLAG-HA* cells with either empty vector (p53-WT) or dominant-negative p53 (p53-DN) grown continuously with 0X Dox, 1X Dox (10ng/ml), or 10X Dox (100ng/l). Plots show mean ± standard deviation for three biological replicates from a single experiment. Statistical significance tested by two-tailed Welch’s t-test with Bonferroni cutoff at a p-value of 0.01 (α = 0.05). No significant differences between p53-WT and p53-DN samples with same Dox treatment. ** = p-value <0.001. N = >1000 nuclei per condition for each replicate. G) Growth curve of MCF10-2A *TetOn-CENPA-FLAG-HA* cells with either empty vector (p53-WT) or dominant-negative p53 (p53-DN) grown continuously with 0X Dox, 1X Dox (10ng/ml), or 10X Dox (100ng/l). Growth curve shows number of population doublings relative to the initial seeding population at Day 0. Dots show mean ± standard deviation for three biological replicates at each time point where cells were counted. Lines show best-fit curves. Similar results were obtained for a second independent experiment. (See Figure S3G for growth curve of the non-inducible parental MCF10-2A cells after Dox exposure).

Importantly, this upregulation is amplified after X-irradiation. Over-representation analysis shows that these genes are mainly involved in cell cycle control, followed by DNA repair (Figure 4E, see also Table S1). We then used cellular assays to investigate the effects of CENP-A overexpression and p53 inactivation on these two pathways. Concerning DNA repair, we did not detect significant changes in either acute DNA damage or rate of DNA repair (Figure S3A-D). However, on a longer timescale, we observed an increase in mitotic stress associated with CENP-A overexpression that was similar in both p53-WT and p53-DN cells (micronuclei in Figure 4F, and CIN in Figure S3E, including numerical and structural aneuploidy by mFISH karyotypes). This is in agreement with previous studies (Athwal et al., 2015; Shrestha et al., 2017). In the p53-WT cells, this mitotic stress was associated with p53 activation, as shown by Western blot (Figure S3F). As for cell cycle control, CENP-A overexpression leads to prolonged growth inhibition that is partially counteracted by p53-DN, an effect that is amplified with increased Dox (Figure 4G, see also Figure S3G for control).

These findings enabled us to discard a major role for DNA damage or repair in the radiosensitivity phenotype. Therefore, based on both the bulk RNA-seq data and cellular assays, we conclude that induced CENP-A overexpression leads to p53-dependent radiosensitivity mainly by impairing cell cycle progression. We then wished to understand how this was operating at the level of individual cells.

### CENP-A overexpression promotes acute cell cycle arrest and senescence in p53-WT cells

To determine the effects of CENP-A overexpression and p53 status on individual cell fate and evolution of the cell population over time, we performed single-cell RNA sequencing (scRNA-seq), a powerful tool for characterizing cell state and identity (Tanay and Regev, 2017). Using our p53-WT and p53-DN MCF10-2A cells, we compared prolonged, continuous CENP-A overexpression (chronic induction), to acute CENP-A overexpression and non-overexpressing conditions (Figure 5A). First, we confirmed the chronic overexpression of CENP-A by Western blot (Figure S4A) and verified that the scRNA-seq data agreed with the bulk RNA-seq data from comparable samples (Figure S4B). Analysis of the single-cell expression profiles revealed that most of the cell-to-cell variability across conditions could be attributed to cell cycle differences (Figure 5B). Indeed, with the scRNA-seq data we could identify distinct sub-populations of cycling and non-cycling cells, and further distinguish G2/M and G1/S cells within the cycling cluster (Figure 5C). Both acute and chronic CENP-A overexpression in the p53-WT cells caused a clear shift in the proportion of cycling to non-cycling cells, a shift that was substantially reduced in the p53-DN cells (Figure 5D-E).

**Figure 5.**
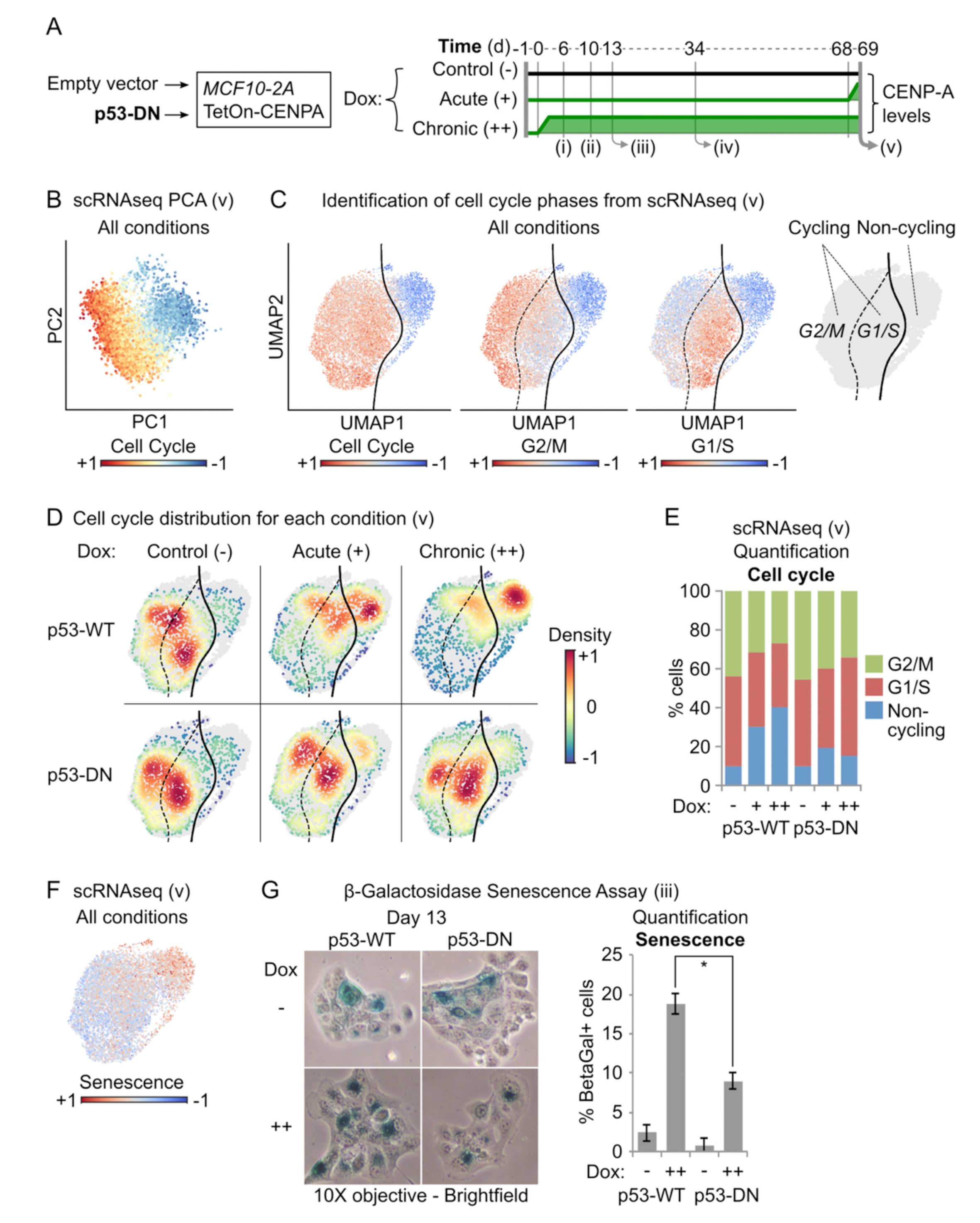
CENP-A overexpression promotes acute cell cycle arrest and senescence in p53-WT cells. A) Scheme delineating conditions tested at various time points (roman numerals), corresponding to Figures 5, 6, S4 and S5, showing relative CENP-A protein levels over time. Note that acute CENP-A overexpression was only tested at time point ‘v’. B) Principal component analysis (PCA) of scRNA-seq experiments at time point ‘v’ in A. All conditions merged. Each dot represents a single cell on the first two principal components (PC1 and PC2), coloured by expression of cell cycle genes (Cell Cycle signature includes all genes from Cyclebase v3.0, *CENPA* excluded). C) Uniform Manifold Approximation and Projection (UMAP) maps of scRNA-seq data from time point ‘v’ in A. All conditions combined. Each dot corresponds to an individual cell projected in a 2D space where the topology reflects global similarities in expression. The colour gradient is proportional to the expression of Cell Cycle, G2/M and G1/S genes (Cyclebase v3.0). Far right plot: boundary between cycling (G2/M or G1/S) and non-cycling cells in UMAP space computed by Support Vector Classification. D) Cell density by condition (colour gradient) in UMAP space for each scRNA-seq experimental condition from time point ‘v’ in A. E) Stacked bar plot showing the % of cells within G2/M, G1/S and non-cycling scRNA-seq clusters at time point ‘v’ in A for each experimental condition. F) UMAP of scRNA-seq data as in C for all conditions merged, coloured by expression gradient of senescence-associated genes (Fridman Senescence Up gene set from MSigDB v6.2). G) Beta-galactosidase senescence assay of samples as in A, at time point ‘iii’. Left: Representative brightfield images after 13 days of growth under the indicated conditions, taken with a 10X objective. Only cells with dark blue staining were considered senescent. Right: quantification of frequency of senescence, showing mean ± standard deviation from two biological replicates of a single experiment. Similar results were obtained in a second independent experiment. * = two-tailed Welch’s t-test comparing p53-WT to p53-DN after chronic CENP-A overexpression; p-value = 0.05.

Interestingly, we found that the non-cycling populations in the scRNA-seq experiments displayed a striking downregulation of the genes corresponding to Cluster 8 in the bulk RNA-seq analysis (Figure S4C), the subset of genes identified as critical for the radiosensitivity phenotype. Given the well-documented role of p53 in senescence (Abbadie et al., 2017; Tonnessen-Murray et al., 2017), we also confirmed that the non-cycling scRNA-seq cluster was consistent with a senescence gene expression signature (Figure 5F). We thus tested the impact of p53 status and prolonged CENP-A overexpression on cell senescence using Beta-galactosidase senescence assays (Figure 5G). p53-WT cells showed on average ∼19% senescence after 13 days of chronic CENP-A overexpression, compared to less than 3% in the non-induced control. Meanwhile, p53-DN cells showed approximately half the levels of senescence (∼9% and ∼1%, respectively). This is consistent with the non-cycling ratios for p53-WT and p53-DN cells in the scRNA-seq data and shows a significant impact on senescence from p53 status. Taken together, our results reveal that CENP-A overexpression causes a clear shift in cell state, promoting acute cell cycle exit and senescence in p53-WT cells that is severely reduced when p53 is defective.

### CENP-A overexpression promotes epithelial-mesenchymal transition in p53-defective cells

Next, we explored if CENP-A overexpression also impacts cell identity in the different p53 contexts. After correcting for cell cycle effects, we re-analyzed the scRNA-seq data for emergent sub-populations deriving from our original epithelial cell lines (MCF10-2A). We thus identified the main clusters of cells (Figure S5A) and characterized the specific markers associated to each cluster (Table S2). We found two subpopulations of epithelial cells with different expression of genes involved in cell metabolism (Figure S5B-C) and a distinct cluster of cells that was negative for epithelial markers. This cluster showed high expression of mesenchymal genes and broad upregulation of genes involved in epithelial-mesenchymal transition (EMT)(Figure 6A-B, see also Table S2). This is consistent with highly advanced EMT or complete mesenchymal transition. The mesenchymal cluster was nearly exclusive to the p53-DN cells (Figure 6C), consistent with previous studies demonstrating that WT p53 counteracts EMT (Brosh et al., 2013; Kim et al., 2013; Pinho et al., 2011; Senoo et al., 2007; Singh et al., 2015; Wang et al., 2009). Remarkably, the highest proportion of mesenchymal cells in the p53-DN condition occurred after chronic CENP-A overexpression. Given that these cells represent a highly advanced stage of EMT, these findings led us to re-explore our cells for earlier EMT signatures by microscopy. We first noticed that large groups of cells with mesenchymal-like characteristics (reduced cell-cell contacts, distorted cell shape) could be observed in the p53-DN cells by simple brightfield microscopy as early as 10 days after continuous CENP-A overexpression (Figure S5E). At Day 34, we immunostained simultaneously for the classic epithelial marker E-cadherin and the mesenchymal marker vimentin (Figure 6D), to identify cells in a broad range of early to late stages of EMT (Pastushenko et al., 2018). The results revealed a major increase in the EMT population after prolonged CENP-A overexpression, which was again exclusive to the p53-DN cells.

**Figure 6.**
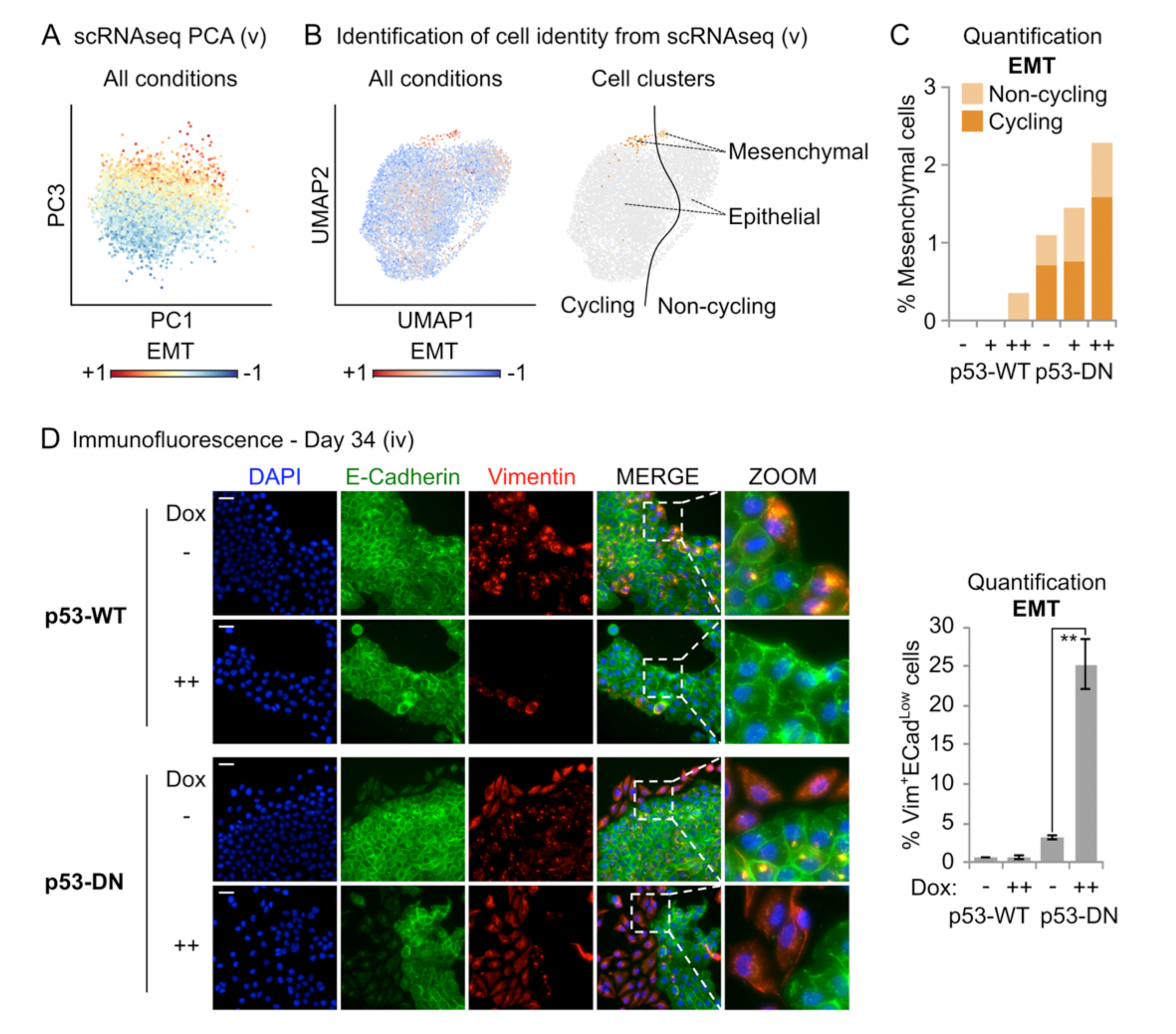
CENP-A overexpression promotes epithelial-mesenchymal transition in p53-defective cells. A) Principal component analysis (PCA) of scRNA-seq experiments at time point ‘v’ in Figure 5A. All conditions merged. Each dot represents a single cell on the first (PC1) and third (PC3) principal components, coloured by expression of epithelial-mesenchymal transition genes (EMT; Hallmark gene set, MSigDB v6.2). Cell-to-cell variability along the third principal component can be explained by differences in expression of EMT genes. B) Classification of epithelial and mesenchymal subpopulations based on Leiden clustering after correcting for cell cycle effects. Left: The colour gradient shows, for each cell, the relative expression trend of genes involved in EMT (EMT; Hallmark gene set, MSigDB v6.2) in UMAP space. Right: Classification of cells as mesenchymal (EMT high cluster, orange) and epithelial (EMT low clusters, grey), divided as either Cycling (dark) or Non-cycling (pale), as determined in Figure 5. C) Stacked bar plots showing the percentage of cells within the mesenchymal cluster per experimental condition. D) Assessment of EMT by immunofluorescence at time point ‘iv’ in Figure 5A. Left: Representative max intensity projections: DAPI (blue), E-cadherin (green, epithelial marker), and Vimentin (red, mesenchymal marker). Scale bars = 40µm. Right: Quantification of the frequency of cells with high Vimentin surrounding the nucleus and low/absent E-cadherin on the cell membrane for each condition. Plots show mean ± standard deviation from three biological replicates. N = >1000 nuclei per condition per replicate. ** = two-tailed Welch’s t-test comparing control p53-DN to p53-DN after chronic CENP-A overexpression; p-value = 0.0002.

Together, these assays reveal that chronic CENP-A overexpression promotes a reprogramming of cell identity in p53-defective cells, corresponding to a striking EMT in the p53-DN cell population.

## Discussion

### CENP-A overexpression drives distinct cell fates depending on p53 status

In this study, we found that CENP-A overexpression promotes radiosensitivity through non-genetic, reversible effects. This is accompanied by major transcriptional reprogramming, involving acute cell cycle shutdown and senescence when p53 is active. However, inactivation of p53 suppresses the cell shutdown response, enabling radio-tolerance.

Strikingly, prolonged CENP-A overexpression also promotes EMT in p53-defective cells. Taken together, our findings demonstrate that CENP-A overexpression promotes two distinct cell fates that depend on p53 status: (i) loss of self-renewal capacity and radiosensitivity in p53-WT cells, and (ii) reprogramming of cell identity through EMT when p53 is defective (Figure 7). Our work reveals an unanticipated link between a centromeric protein and genome reprogramming with clear implications for tumour evolution. These findings open up exciting avenues for future research and have broad implications for cancer treatment.

**Figure 7.**
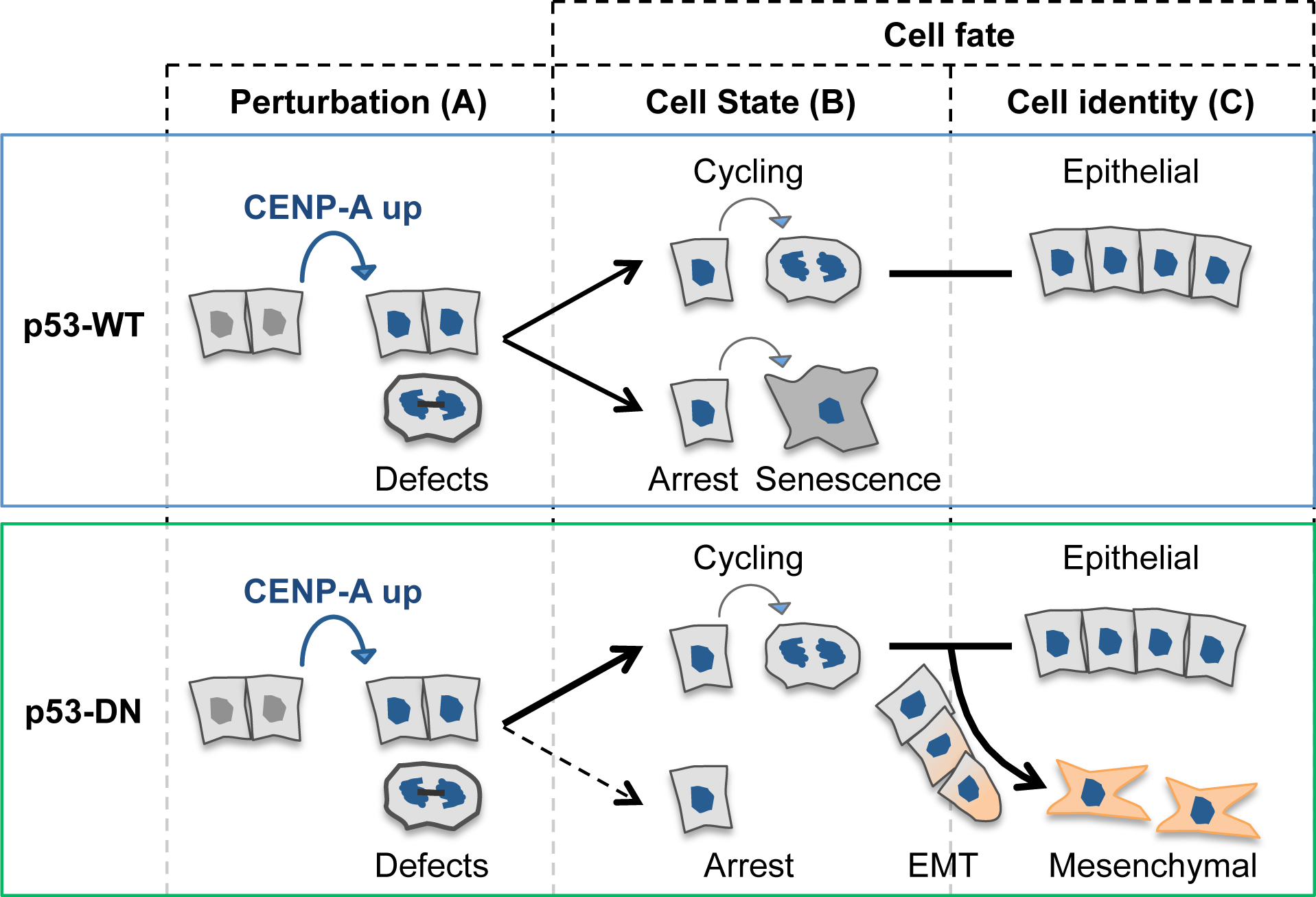
CENP-A overexpression drives distinct cell fates depending on p53 status. CENP-A overexpression reprograms cell fate with distinct effects on cell state and cell identity that depend on p53 status. Perturbation (A) by CENP-A overexpression (in blue) induces mitotic defects in both p53-wild type (WT, top panel blue) and p53-defective cells (DN, dominant negative, bottom panel green). These defects provoke distinct cell fate decisions according to p53 status, impacting cell state (B) or identity (C). When p53 is functional, cell state shifts towards acute cell cycle arrest and senescence, reducing cell self-renewal. Additional stress, like DNA damage from X-irradiation, amplifies this response, resulting in radiosensitivity. Furthermore, functional p53 ensures the preservation of epithelial identity. In contrast, when p53 is defective, the cells evade arrest and continue cycling, allowing CENP-A overexpression to promote epithelial-mesenchymal transition (EMT).

### Induced CENP-A overexpression alters cell state and global transcription: implications for cancer treatment

Our work shows that at the time scale of one cell division or less, induced CENP-A overexpression in MCF10-2A cells results in major transcriptional reprogramming across the genome. Switching p53 status from wild type to defective counteracted a subset of these transcriptional changes, corresponding to cell cycle genes. But, while our scRNA-seq analyses revealed that these effects on cell cycle genes at the population level can be explained by changes in the proportion of arresting cells, how CENP-A overexpression leads to such broad and rapid changes to gene expression across the genome remains an open question. Given that CENP-A overexpression results in its mislocalization and incorporation into chromatin across the chromosome arms, including numerous genic loci (Athwal et al., 2015; Lacoste et al., 2014; Nechemia-Arbely et al., 2019; Nye et al., 2018), we envisage two main scenarios for how CENP-A impacts global transcription: (i) the changes reflect a response to defects induced by CENP-A overexpression and/or, (ii) CENP-A incorporation into chromatin directly affects transcription at the sites where it is mislocalized. The idea that ectopic CENP-A could directly affect transcription at genic loci has recently been proposed (Saha et al., 2020), but remains to be formally assessed. Understanding the underlying mechanisms of these effects could provide important insights into alternative functions of CENP-A and its impact beyond the centromere. On a longer time-scale, our scRNA-seq results revealed that prolonged CENP-A overexpression promotes chronic cell cycle arrest in a major proportion of p53-WT cells. Thus, our findings support a model where CENP-A overexpression induces mitotic defects and increases CIN (Shrestha et al., 2017), which, in turn, lead to the activation of p53 and senescence (Hinchcliffe et al., 2016; Soto et al., 2017). In this way, CENP-A overexpression causes a p53-dependent loss of self-renewal capacity. Additional cell stress from DNA damaging agents, like X-irradiation, would add to the mitotic stress associated with CENP-A overexpression. This would then amplify the p53-dependent shutdown of self-renewal, promoting radiosensitivity. Thus, our findings suggest that differences in p53 status may be able to explain why high CENP-A levels are associated with both sensitivity (McGovern et al., 2012; Sun et al., 2016; Zhang et al., 2016) and resistance (Gu et al., 2014) to cancer treatments in different patient cohorts. This could have important implications for patient prognosis and treatment strategies. Radiosensitivity associated with high CENP-A levels in p53-WT cancers could represent an opportunity to stratify patients or minimize radiotherapy dose for equal therapeutic outcome with reduced side-effects. Furthermore, alternative treatment options for patients with non-functional p53 could also be considered, as drugs combatting mutant p53 (including LOF mutations) are currently in development (Blandino and Di Agostino, 2018; Bykov et al., 2018). A number of these drugs are now in clinical trials, including direct re-activators of mutant p53 (e.g.PRIMA-1^MET^/APR-246) and drugs designed to return functionality to WT p53 by inhibiting its upstream regulators (e.g. MDM2/X inhibitors)(Sanz et al., 2019). Here, we showed that transiently reactivating p53 in CENP-A overexpressing HeLa cells was able to significantly increase sensitivity to X-irradiation. Thus, by returning functional p53 status to tumours with high CENP-A, not only could this induce the anti-tumoural effects of activating p53 alone, but it may also promote radiosensitivity due to the effects of CENP-A overexpression on cell state.

### Prolonged CENP-A overexpression alters cell identity: implications of EMT

One of the most surprising findings from our study was the striking emergence of EMT in p53-defective cells after prolonged CENP-A overexpression. EMT is a multi-stage process where epithelial cells, characterized by strong cell-cell junctions and apical-basal polarity, undergo a series of changes to gene expression and morphology to gain a mesenchymal phenotype, including a reorganized cytoskeleton, altered cell shape, and increased cell motility (Lamouille et al., 2014). While this transition from epithelial to mesenchymal is an essential process in mammalian development, it is also considered a key step in the development of invasion and metastasis in epithelial cancers, with impacts on proliferation, cell plasticity, stemness, and therapeutic resistance (Pastushenko and Blanpain, 2019; Puisieux et al., 2018). CENP-A overexpression has previously been linked to invasion and metastasis in human patients (Gu et al., 2014; Ma et al., 2003; McGovern et al., 2012; Saha et al., 2020; Sun et al., 2016; Zhang et al., 2016), but whether this could involve promotion of EMT was not known. Thus, the link with EMT provides a mechanism by which CENP-A overexpression could drive these outcomes in patient tumours that lack functional p53.

Evidence linking EMT to the direct inhibition of senescence (Ansieau et al., 2008; Liu et al., 2008), promotion of radioresistance (de Jong et al., 2015; Steinbichler et al., 2018; Theys et al., 2011), and cancer stemness (Shibue and Weinberg, 2017), imply that EMT may also play an important role in the evasion of senescence, radio-tolerance, and increased clonogenic capacity that we observed in our p53-defective cells. Interestingly, the effect of CENP-A overexpression on EMT also suggests that high CENP-A levels could promote stemness, depending on p53 status. Indeed, human pluripotent stem cells naturally overexpress CENP-A (Ambartsumyan et al., 2010; Milagre et al., 2020), and maintain p53 in an inactive state through post-translational regulation (Jain and Barton, 2018). Intriguingly,recent work in Drosophila showed that intestinal stem cells preferentially retain pre-existing CENP-A during asymmetric divisions, suggesting that CENP-A nucleosomes may epigenetically mark stem cell identity in this system (García del Arco et al., 2018). Thus, in addition to its effect on EMT, how CENP-A overexpression contributes to stem cell renewal and pluripotency in human cells will be an important avenue for future research. Importantly, as with the effect of CENP-A overexpression on global transcription, which underlying mechanism is at play remains to be deciphered. Since the initiation of EMT is associated with several of the cell stresses that result from mitotic defects, including genotoxic stress (Wu et al., 2013), replication stress (McGrail et al., 2018), and metabolic stress (Cha et al., 2015), amongst others (Chen et al., 2017; Jiang et al., 2017; Marcucci and Rumio, 2018; Shah and Beverly, 2015), they are likely important in the process. Interestingly, a recent report by Gomes et al demonstrated that the perturbation of the histone H3 variants, H3.1/2 and H3.3, promotes or represses EMT (Gomes et al., 2019). In particular, knockdown of the H3.1/2-dedicated chaperone CAF-1 caused widespread opening of chromatin, with increased incorporation of H3.3 at the promoters of EMT-inducing transcription factors (e.g. *ZEB1*, *SNAI1*, and *SOX9*), which induced EMT and increased cell migration and invasion.

Previously, we found that overexpressed CENP-A hijacks the H3.3-dedicated chaperone Daxx (Lacoste et al., 2014). This results in the mis-incorporation of CENP-A-containing nucleosomes into regions of high histone turnover normally enriched for H3.3. Thus, CENP-A overexpression could have a direct impact on EMT, and transcription in general, through its perturbation of other H3 variants and their dedicated histone chaperones. Deciphering the mechanisms that link CENP-A, EMT, and possibly stemness, will expand our understanding of the direct and indirect molecular consequences of CENP-A overexpression and its impact on tumour evolution.

In conclusion, the interplay of CENP-A overexpression and p53 status alters cell fate, with distinct implications for the role of CENP-A in therapeutic sensitivity, resistance and metastasis.

## Acknowledgements

We sincerely thank Jean-Pierre Quivy and Leanne De Koning for critical reading of the manuscript. We also thank Patricia Le Baccon and the Cell and Tissue Imaging Platform – PICT-IBiSA (member of France–Bioimaging – ANR-10-INBS-04) of the UMR3664 of Institut Curie for help with light microscopy, as well as Sophie Heinrich and the Experimental Radiobiology platform of Institut Curie for help with X-irradiation. High-throughput sequencing was performed by the ICGex NGS platform of the Institut Curie supported by the grants ANR-10-EQPX-03 (Equipex) and ANR-10-INBS-09-08 (France Génomique Consortium) from the Agence Nationale de la Recherche (“Investissements d’Avenir” program), by the Canceropole Ile-de-France and by the SiRIC-Curie program - SiRIC Grant « INCa-DGOS- 4654. DF receives salary support from the Centre Nationale de Recherche Scientifique (CNRS). MD receives salary support from the City of Paris via Emergence(s) 2018 of DF. AG, DJ, KP, LB, RPL and GA were supported by la Ligue Nationale contre le Cancer (Equipe labellisée Ligue), Labex DEEP (ANR-11-LABX-0044_DEEP, ANR-10-IDEX- 0001-02), PSL, ERC-2015-ADG-694694 ChromADICT and ANR-16-CE12-0024 CHIFT. Funding for RPL provided by Horizon 2020 Marie Skłodowska-Curie Actions Initial Training Network “EpiSyStem” (grant number 765966). Individual funding was also provided to DJ from la Fondation ARC pour la recherche sur le cancer (“Aides individuelles” 3 years, post- doc), and to AG from the Horizon 2020 Framework Programme for Research and Innovation (H2020 Marie Skłodowska-Curie Actions grant agreement 798106 “REPLICHROM4D”).

## Author Contributions

Conceptualization, DJ and GA; Methodology, DJ, KP and GA; Software, AG; Validation, AG, DJ, KP, LB, and RPL; Formal Analysis, DJ and AG; Investigation, AG, DJ, KP, LB, MD and RPL; Resources, DF and GA; Data Curation, AG; Writing – Original Draft, DJ, AG and GA; Writing – Review & Editing, DJ, KP, LB, RPL, MD, DF and GA; Visualization, AG, DJ, KP, MD and GA; Supervision, DF, KP and GA; Project Administration, GA; Funding Acquisition, GA.

## Materials and Methods

### Cell lines

| Cell line | p53 status* | Cell type | Construct | Comments |
| --- | --- | --- | --- | --- |
| *MCF10-2A* | Functional wild type | Immortal non-tumoural (fibrocystic disease) breast epithelial | *TetOn-CENP-A-FLAG-HA* (lentiviral) |  |
| *HeLa S3* | Non-functional wild type: HPV18 inactivated | Cervical cancer | *TetOn-CENP-A-FLAG-HA* (lentiviral) |  |
| *T47D* | LOF p53 mutation at L194F | Metastatic-derived ductal carcinoma, breast | *TetOn-CENP-A-FLAG-HA* (lentiviral) | HER<sup>-</sup> ER<sup>+</sup> PR<sup>+</sup> Basal A |
| *HCC1954* | LOF p53 mutation at Y163C | Invasive ductal carcinoma, breast | *TetOn-CENP-A-FLAG-HA* (lentiviral) | HER<sup>+</sup> ER<sup>-</sup> PR<sup>-</sup> luminal |
| *DLD1* | One functionally silent allele of p53 & LOF mutation at S241F | Colon cancer | *TetOn-CENPA-YFP-AID* (FlpIn) | Kind gift from D. Fachinetti |

\*p53 status for missense mutations from the TP53 database (Bouaoun et al., 2016).

### Cell culture and CENP-A overexpression

HeLa S3, DMEM 10% Fetal Cow Serum (FCS) Pen/Strep 1%; MCF10-2A, 1:1 DMEM:Ham’s F12, 20 ng/ml epidermal growth factor, 100 ng/ml cholera toxin, 0.01mg/ml insulin and 500 ng/ml hydrocortisone, 5% horse serum; T47D and HCC1954, RPMI 10% FCS Pen/Strep 1%; DLD-1 FRT Fbox (+OSTR1-Myc9)+CENP-A-YFP-AID (resistant to hygromycin, puromycin and blasticidine), DMEM 10% Tet-free FCS (Thermofisher Scientific) Pen/Strep 1%. Base media from Thermofisher Scientific. All cell lines tested and confirmed to be mycoplasma free using the Mycoplasma PCR ELISA kit (Sigma). CENP-A overexpression was induced by the addition of Dox to typical growth media at 10ng/ml (considered 1X), unless otherwise indicated. To maintain overexpression, media and Dox was replaced every one to three days. When applicable, the media in the non-induced control was replaced in parallel without Dox. For reversal of CENP-A overexpression, we removed media containing Dox and washed two times with warm PBS then replaced the media without Dox.

### Generation of *TetOn-CENPA-FLAG-HA* cell lines

To obtain an inducible CENP-A overexpression system, we sub-cloned *CENPA-FLAG-HA* from the eCENP-A plasmid (Lacoste et al., 2014) by PCR (Forward primer containing EcoRI site: *CACAGTTA<u>GAATTC</u>**ATG**GGCCCGCGCCG*; Reverse primer containing AgeI site:

*ATCGAATC<u>ACCGGT</u>**CTA**GGCGTAGTCGGGCACGT*) and inserted it into the multiple cloning site of the pLVX-TetOne (ClonTech) plasmid by restriction enzyme digestion with EcoRI and AgeI, followed by ligation. To confirm proper integration and lack of mutation in the *CENPA* gene, we then sequenced the *CENPA-FLAG-HA* pLVX-TetOne plasmid around the insertion site. To encapsulate the construct for transduction we co-transfected Lenti-X 293T cells (ClonTech) with 10µg *CENPA-FLAG-HA* pLVX-TetOne and packaging vectors 7.5µg psPAX2 and 2.5µg pMD2.G using PolyPlus JetPrime transfection reagent according to manufacturer instructions. Media was replaced 4h after transfection. For infection and stable integration of the construct without selection, we filtered (0.45µm) supernatant from the transfected cell flask at 24h and 48h post-transfection, added 8µg/ml of polybrene to the filtered virus-containing media, and added the media directly to the target cell lines. We removed the virus-containing media after 48h and grew the cells for at least three passages without virus. Then we tested the polyclonal cell lines for the induction of *CENPA-FLAG-HA* by Western and IF of CENP-A and HA following 24h of treatment with 80ng/ml of doxycycline. We isolated clonal cell lines by serial dilution to single cells. Then tested 10-20 colonies arising from single cells for *CENPA-FLAG-HA* induction, as before. We selected cell lines for further experimentation if they showed a clear homogenous CENP-A and HA increase by IF in ∼100% of cells after doxycycline treatment with no detectable background of HA signal by Western or IF when no doxycycline was added.

### Change of p53 status

For the p53-WT and p53-DN MCF10-2A cells, we transduced empty vector pWZL Hygro (Scott Lowe, Addgene plasmid #18750) or vector containing the dominant negative *TP53* construct (pBABE-hygro p53 DD, Bob Weinberg, Addgene plasmid #9058) into the clonal MCF10-2A *TetOn-CENPA-FLAG-HA* cell line by lentiviral transduction, as above. Cells were selected with hygromycin for >14 days and then passaged in typical media. For heat shock of HeLa S3 *TetOn-CENPA-FLAG-HA cells*, we performed a 1h 42°C heat shock on non-induced HeLa *TetOn-CENPA-FLAG-HA* cells and examined total cell extracts by Western blotting at various time points following heat shock. Two parallel sets of cells were grown under normal conditions at 37°C. One set was moved to a 42°C incubator while the other remained at 37°C as a control. After 1h, we harvested cells for total cell extraction from each condition (time 0h), then returned both sets to 37°C. We then harvested cells for total cell extraction at the indicated times after heat shock (1h, 2h, 3h, 4h, or 6h: Time after HS).

### Total cell extracts

We harvested cells by trypsin, counted cells by Beckman Automated Cell Counter, and spun down at 300g, 5 min, washed 1X in PBS, spun again, aspirated PBS and froze pellets at - 20°C. Cell pellets were incubated for 15min at room temperature with 300µl per 1 million cells of 1.2X LDS Sample Buffer (NuPAGE) containing 1.2X Sample Reducing Agent (NuPAGE) and 125kU/ml Pierce Universal Nuclease Buffer for Cell Lysis, then heated at 95°C for 10min, vortexed, spun down, and cooled briefly on ice prior to Western blotting.

### Western blotting

Total cell extracts were loaded at two to four different concentrations, as indicated, onto pre-made NuPAGE Bis-Tris 4-12% gradient gels in an XCell 4 Sure-Lock SDS-PAGE chamber with 1X NuPAGE MES SDS Running Buffer, with a PageRuler (Thermofisher scientific) molecular weight marker. Gels were run at 130-150v for 1h-1h15min. We transferred protein to a 0.2µm nitrocellulose membrane by BioRad Trans-Blot Turbo, mixed-molecular weight setting, or semi-dry transfer (BioRad, 15v, 1.5h). Membranes were stained with Pierce reversible protein stain to detect bulk protein and assess quality of transfer, then cut, blocked in 5% milk-PBST (0.1% Tween-20) for 1h, RT, and incubated overnight with rocking at 4°C with primary antibodies in 5% milk-PBST containing 0.02% sodium azide. Membranes were washed 3X 10min in PBST, then incubated 1h (RT) with Horse Radish Peroxidase (HRP) secondary antibodies in 5% milk-PBST, washed 3X 10 min in PBST and exposed with SuperSignal West Pico Chemiluminescent Reagent or, for high sensitivity exposures, SuperSignal West Femto Maximum Sensitivity Reagent, and imaged using a ChemiDoc Touch system and ImageLab software. Primary antibodies: CENP-A 1:500 (2186 Cell Signaling Technology (CST)), H4-pan 1:2500 (05-858, Sigma-Aldrich), HJURP 1:300 (HPA008436 Sigma-Aldrich), γTubulin 1:10000 (T5326 Sigma-Aldrich), p53 (1C12) 1:1000 (2524 CST), p21 1:500 (556431 BD Pharmingen), phospho-p53 (Ser15) 1:500 (9284 CST).

Secondary antibodies: Jackson ImmunoResearch, 1:10000, donkey anti-mouse or donkey anti-rabbit.

### Immunofluorescence (IF)

Glass cover slips were coated with collagen (1µg/ml) + fibronectin (1µg/ml) in PBS, 30min, RT, washed in PBS and added to culture dishes prior to cell seeding. After at least 24h of growth, coverslips were washed 3X with PBS, fixed with 2% PFA, 20min, RT, washed 3X PBS, incubated 5 min with 0.3% Triton-X 100 in PBS, washed 3X PBS, blocked in filter-sterilized 5% BSA-PBST (0.1% Tween-20) for 20-30min, RT, and incubated 1-1.5h, RT, with primary antibodies in 5% BSA-PBST. Coverslips were then washed 3X 5min in 5% BSA-PBST, and incubated 30min with secondary antibody in 5% BSA-PBST, RT, in darkness.

DAPI was added directly to secondary antibody solution (final concentration 1/4000) and incubated in darkness, RT, 5min, followed by 3X PBS wash. Coverslips were inverted onto microscope slides with ∼10µl of VectaShield and imaged with a Zeiss Axiovert Z1 microscope or Inverted Widefield Deltavision Core Microscope (Applied Precision) with CoolSNAP HQ2 camera. Primary antibodies: CENP-A (3-19) 1:300 (ADI-KAM-CC006-E Enzo Life Sciences), E-cadherin (24E10) 1:200 (3195S CST), γH2AX 1:250 (2577 CST), Vimentin (N-term) 1:100 (5741S Progen). Secondary antibodies: 1:1000 Alexa Fluor donkey anti-mouse IgG (H+L) 488, 1:1000 Alexa Fluor goat anti-rabbit IgG (H+L) 488 or 594, 1:1000 Alexa Fluor goat anti-Guinea Pig IgG (H+L) 594.

### Colony Formation Assays (CFAs)

At Day -1, cells are trypsinized and diluted to single-cell level (300-600 cells per well, in triplicate) in 6-well plates. At Day 0, one set of cells is irradiated by CIXD Dual Irradiator or Philips X-ray tube X-ray generator (4gy, unless otherwise indicated) while a control set remains un-irradiated (0gy). When the cells have had sufficient time to form visible colonies, depending on their typical speed of growth (7-14 days), they are washed gently with PBS, stained with 1% Crystal Violet 20% Ethanol solution for 15 min, washed with water, allowed to dry, scanned and counted by eye from the acquired images with the counter blinded to the conditions. Unless otherwise indicated, ratio of surviving colonies is calculated as the number of colonies formed after 4gy of X-irradiation divided by the number of colonies formed when not irradiated. Cells are considered sensitized to irradiation by CENP-A overexpression when the CENP-A overexpression condition has a significantly lower survival ratio compared to the non-induced control. Cells are considered tolerant to CENP-A overexpression if the survival ratio is not significantly affected by the CENP-A overexpression.

### Bulk RNA-seq

Sample preparation: We grew MCF10-2A *TetOn-CENPA-FLAG-HA* cells expressing either empty vector (p53-WT) or dominant-negative p53 (p53-DN) with 0X Dox (no Dox), 1X Dox (10ng/ml), or 10X Dox (100ng/m) for 24h. At time 0, we irradiated one set of cells by X-ray generator (4gy) while a control set remained un-irradiated (0gy). 6h later, we extracted RNA for RNA-seq. All conditions tested in duplicate. We extracted RNA directly from culture dishes using the Qiagen RNeasy Mini kit according to the manufacturer’s instructions and confirmed the quality and quantity of the RNA by TapeStation and NanoDrop. mRNA library preparation and sequencing were performed by the Next Generation Sequencing (NGS) platform, Institut Curie, using the Illumina TruSeq high input RNA kit and NovaSeq 6000 sequencer. Library preparation and sequencing of irradiated and non-irradiated samples were performed on different dates, with a single repeat of one sample included for batch correction.

### Bulk RNA-seq analysis

Reads were aligned to the human reference genome (GRCh38 assembly) based on Ensembl gene annotation (release 95) with hisat2 (version 2.1.0; Kim et al., 2019), run in paired-end mode with default parameters. Gene-level counts were computed from primary alignments with MAPQ > 2 using featureCounts (Subread package version 1.6.3; Liao et al., 2014) in paired-end mode with the -s 2 option for reverse-stranded libraries. Raw counts were normalized for differences in library size (counts per million) and across samples (via trimmed mean of M-values normalization, TMM) using edgeR (version 3.28.0; Robinson et al., 2010). Mean-centered TMM-normalized counts were used for PCA and hierarchical clustering analyses via Ward’s variance minimization method. Differential expression analyses were performed with edgeR (McCarthy et al., 2012). We assessed the stand-alone and combined effect of each treatment (CENP-A overexpression at increasing Dox concentrations, p53-DN, and X-irradiation) by fitting a quasi-likelihood negative binomial generalized log-linear model (GLM). A batch coefficient was also included to account for potential batch effects. The effect of each treatment, both stand-alone and combined, was evaluated via quasi-likelihood F-test on the respective GLM coefficient, followed by multiple testing correction via the Benjamini-Hochberg method (Table S1). A false discovery rate (FDR) cut-off of 0.05 was used to identify differentially expressed genes (DEGs). For proportional Venn diagrams, we only considered DEGs that change with CENP-A overexpression, p53 status or X-irradiation, since we could not detect significant interaction effects for most treatment combinations (Table S1). For hierarchical clustering, we tested the combined effect of all coefficients, excluding batch, and included any DEG showing significant differences relative to baseline at 0.05 FDR. Functional enrichment analyses for KEGG pathways were carried out using the WebGestaltR package (version 0.4.2; Liao et al., 2019). Gene set enrichment analyses (GSEA) for CENP-A effects were performed on all expressed genes ranked by fold change and p-value upon overexpression i.e. log _2_ fold change * - log _10_ p-value of the Dox coefficient. Pathways associated to specific gene clusters were identified by over-representation analysis (ORA) after hierarchical clustering of all DEGs via Ward’s method. Coordinated changes in expression within DEG clusters were evaluated by assessing the distribution of mean-centered expression values (averaged per condition) for all genes in a given cluster. Bulk RNA-seq analyses were carried out with custom Python scripts. pandas (version 0.24.2), NumPy (version 1.16.2), SciPy (version 1.3.1), and scikit-learn (version 0.21.3) libraries were used for data manipulation, statistical analysis and unsupervised learning. matplotlib (version 2.2.4), matplotlib-venn (version 0.11.5) and seaborn (version 0.9.0) were used for plotting and statistical data visualization. R packages were imported into Python using rpy2 (version 2.8.4).

### Comet assays

Prior to irradiation, cells were harvested and submerged into media containing low-melting agarose on ice. When hardened, cells were exposed to the indicated dose of γ-irradiation (GSR D1 gamma ray). Either immediately, or at the indicated times post-irradiation, the cells in the gel were submerged in alkaline solution and run by gel electrophoresis to form comets to visualize the levels of DNA damage. Comet assays were performed by the RadExP platform, Institut Curie, according to the Trevigen Single Cell Electrophoresis Assay alkaline comet protocol. Automatic counting and measuring of comets on Trevigen slides (two wells used for each condition, per experiment), imaged using the Metafer system with the “comet” module from MetaSystems.

### Micronuclei

We seeded cells into dishes containing collagen/fibronectin-coated glass coverslips 48h prior to fixation, added indicated concentrations of Dox and allowed the cells to grow for 24h (Day 1) or passed cells and replaced Dox every 2-3 days until Day 8. Cells were fixed, prepared for IF, and nuclei were imaged with a Zeiss Z1 inverted microscope using a 10X objective.

We counted micronuclei using a custom ImageJ macro with at least three fields and >1000 nuclei counted per condition.

### Multicolor fluorescence in situ hybridization (mFISH) karyotyping

Cells were grown with or without 10 ng/ml Dox (as in the growth curve assay) for 15 days, then treated with colcemid (100 ng/ml, Roche) for 1.5 hours and prepared as previously described for mFISH (MetaSystems) staining (Trott et al., 2017). Briefly, mitotic cells were collected by mitotic shake-off after a short trypsin treatment and centrifuged at 1000 rpm for 10 min. Cell pellets were resuspended in 75 mM KCl and incubated for 15 min in a 37°C waterbath. Carnoy fixative solution (methanol/acetic acid, 3:1) was prepared and 1:10 volume added on the cells, before centrifugation at 1000 rpm for 15 min. Cells were then fixed 30 min at room temperature in the Carnoy solution, centrifuged and washed once more with fixative. Minimum volume of fixative was left to resuspend the pellet and cells were dropped onto clean glass slides. Multicolor FISH (mFISH) staining was performed following manufacturer’s instructions (MetaSystems). The Metafer imaging platform (MetaSystems) and the Isis software were used for automated acquisition of the chromosome spread and mFISH image analysis. Chromosome rearrangements and specific chromosome counts for each spread were assessed and counted by eye from the automated chromosome spread images, with the researchers blinded to the conditions. Losses and gains of chromosomes per cell were calculated as the sum of the difference from the mean for each chromosome of the p53-WT no Dox control. New chromosome rearrangements per cell were determined as the total number of structural chromosomal anomalies observed per spread, excluding the ones that were common to all p53-WT no Dox spreads. Potential acrocentric fusions were not considered in our study.

### Growth curve / Proliferation assays

Cells that had never been exposed to Dox were trypsinized, counted using a Beckman Automated Cell Counter, and seeded into nine culture dishes with equivalent cell numbers (Corresponds to MCF10-2A non-inducible control, plus the p53-WT and p53-DN TetOn- CENP-A-FLAG-HA cell lines, at three Dox concentrations each, all grown in parallel). The next day (Day 0), Dox was added in triplicate at the indicated concentrations (0, 10, or 100ng/ml), corresponding to 0X, 1X or 10X Dox. At Day 1, we trypsinized and counted the cells from all conditions, and seeded equivalent numbers of cells into fresh media. We immediately added the corresponding concentrations of Dox to the culture dishes. From then on, every 2-6 days cells were trypsinized, counted and re-plated, as plotted, with media containing fresh Dox of corresponding concentrations replaced every 2-3 days. Biological replicates were maintained separately throughout the entirety of the experiment.

### Single-cell RNA-seq

Sample preparation: We grew MCF10-2A *TetOn-CENPA-FLAG-HA* cells expressing either empty vector (p53-WT) or dominant-negative p53 (p53-DN) without Dox or with continuous exposure to Dox (10ng/ml, ++, chronic) for 69 days in parallel. Cells in the no Dox condition were split into two dishes at day 67 and Dox was added to one set on day 68 for 24h of CENP-A overexpression (10ng/ml, +, acute) while the other remained without Dox (-, control). We prepared samples in singlet according to the 10x Genomics Sample Preparation Demonstrated Protocol. In brief, we harvested cells by trypsinization, resuspended in typical media and mixed thoroughly by pipette, passed cells through a 40µm cell strainer, and counted cells by Beckman Automated Cell Counter. We resuspended cells in 1x PBS + 0.04% BSA, mixed, centrifuged gently, aspired supernatant and repeated wash two times. We passed cells through another cell strainer (40µm), counted again and diluted to a final concentration between 700 and 1200 cells/µl. We then proceeded directly to GEM generation and barcoding at the NGS platform, Institut Curie, using the Single-cell 3’ Reagent Kits v2 protocol with a targeted cell recovery of 2000 cells per sample, followed by post GEM-RT cleanup and cDNA amplification, then 3’ Gene Expression Library Construction, according to the manufacturer’s instructions. Sequencing was performed by the NGS platform with a NovaSeq 6000 sequencer.

### Single-cell RNA-seq analysis

Reads were pseudoaligned to Ensembl transcripts (GRCh38, release 95) using kallisto (version 0.46.0; Bray et al., 2016) with the -x 10x2 option for Chromium Single Cell 3’ v2 chemistry. Count matrices were generated from sorted BUS files using bustools (version 0.39.2; Melsted et al., 2019), after barcode correction with the 10x v2 whitelist. For each sample, we selected cells via distance-based estimation of the knee in the cumulative distribution of distinct UMIs per barcode (unique molecular identifiers) using UMI-tools (version 1.0.0; Smith et al., 2017). Sample matrices were thus converted to AnnData objects and concatenated for further quality control (QC) and analysis with SCANPY (version 1.4.6; Wolf et al., 2018). From the merged count matrix, we filtered out cells with abnormal levels of mitochondrial RNA, after adjusting for depth and total number of detected genes. Outliers were detected by covariance estimation (elliptic envelope with 5% contamination) using the number of genes, total counts (log-transformed) and mitochondrial counts (log-transformed) as features for outlier detection. Genes detected in less than 100 cells (after filtering) were also excluded. For comparison of scRNA-seq and bulk RNA-seq results, raw counts from matched samples were pooled across all cells, TMM-normalized and mean-centered.

For single-cell analyses, raw counts were normalized per cell, log-transformed, and adjusted for differences in sequencing depth via linear regression (using log-transformed total counts per cell). PCA was performed on the top 3000 highly variable genes (HVGs) after scaling.

We computed a neighborhood graph using the first 20 principal components, adjusting for technical differences by batch alignment with BBKNN (version 1.3.9; Polanski et al., 2019). Uniform Manifold Approximation and Projection (UMAP) was used to visualize cell-to-cell variation in a low-dimensional (2D) space. Cell subpopulations were detected by Leiden clustering (at 0.5 resolution). Epithelial and mesenchymal cells were separately identified with the same procedure, but normalized counts were also adjusted for cell cycle differences (using the G1/S and G2/M signature) prior to scaling and PCA. For each cluster, genes were ranked by one-vs-all logistic regression to identify the top markers (See Table S2). Boundaries among clusters were computed via Support Vector Classification with a 3^rd^ degree polynomial kernel. Gene sets for computing expression signatures were retrieved from Cyclebase v3.0 (Cell Cycle, G1/S and G2/M) (Santos et al., 2015) or MSigDB v6.2 (Hallmark gene sets: Epithelial Mesenchymal Transition, Oxidative Phosphorylation, Fatty Acid Metabolism and Glycolysis; Curated gene sets: Fridman Senescence Up) (Liberzon et al., 2015). The scores (colour gradients) are a measure of the average expression of the given set of genes, relative to a set of randomly sampled genes at comparable expression levels. Log-transformed normalized counts (prior to scaling) were used for the calculation. The score is then rescaled across cells from -1 (lowest) to +1 (highest). Singe-cell RNA-seq analyses were carried out with custom Python scripts using SCANPY (version 1.4.6).

BBKNN (version 1.3.9), UMI-tools (version 1.0.0), pandas (version 0.25.0), NumPy (version 1.17.0), scikit-learn (version 0.21.3), matplotlib (version 3.3.0), and seaborn (version 0.9.0).

### Beta-galactosidase senescence assay

We tested samples in duplicate for senescence using the Cell Signaling Technologies Senescence β-Galactosidase Staining Kit, according to the manufacturer’s instructions. After 24h with the stain, we imaged at least three fields per sample by brightfield microscopy using a 10X objective on a Nikon Eclipse TS100 microscope. Cells were counted manually from the acquired images with the counter blinded to the conditions. Only cells with dark blue staining were considered senescent.

