## Supplemental Figures for "CENP-A overexpression drives distinct cell fates depending on p53 status"

Supplemental Information

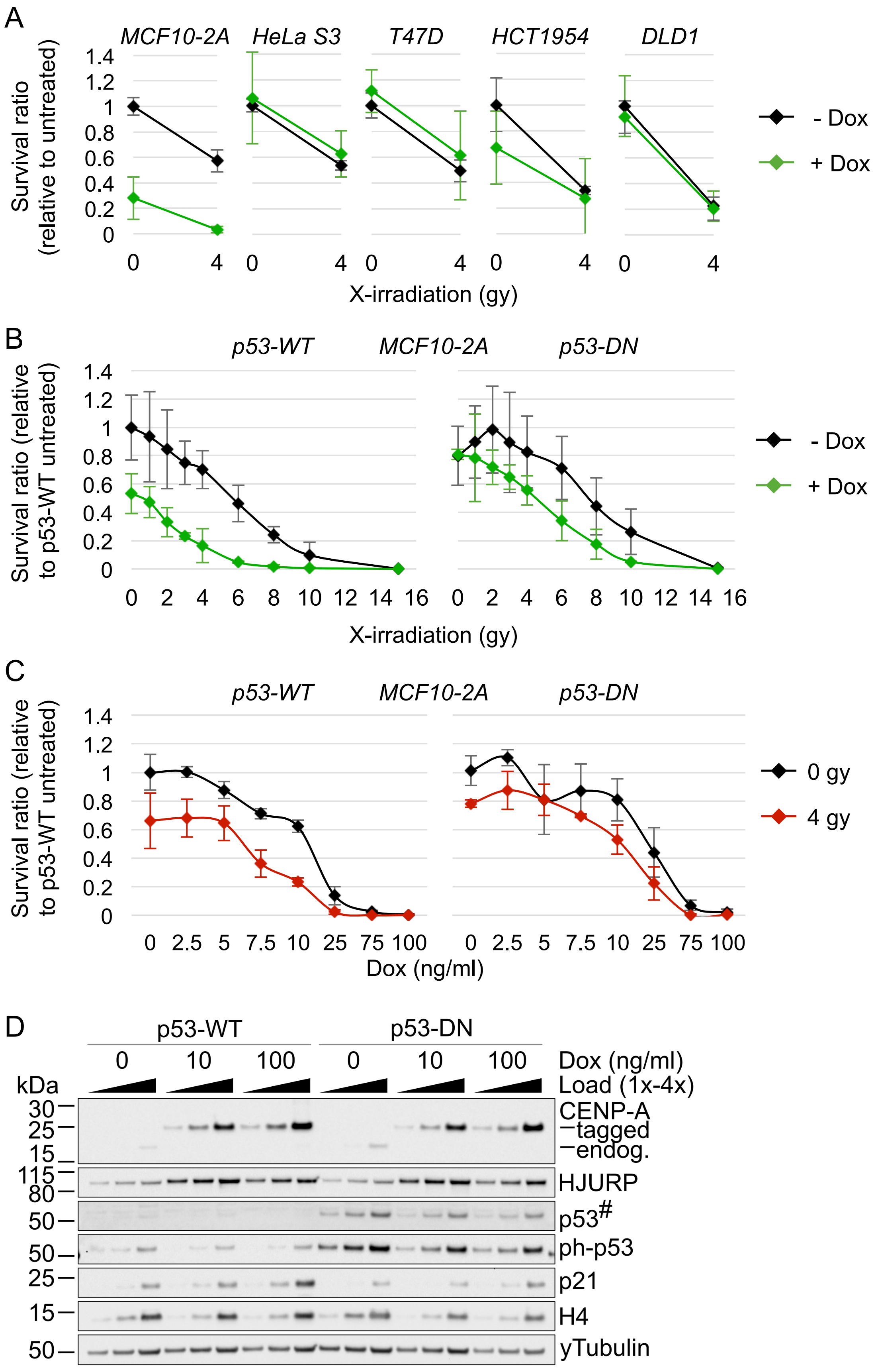


Figure S1

**Figure S1 – Related to Figures 2, 3, 4 and S3. Defined range of p53-dependent radiosensitivity in MCF10-2A cells**

A) Line plots of colony formation assay (CFA) results from Figure 2B, except showing survival ratios relative to untreated condition (0gy, no Dox) for each cell line. Plots show mean ± 95% confidence interval.

B) CENP-A overexpression causes p53-dependent X-irradiation sensitivity at a defined range of X-irradiation doses. CFAs as in Figure 2D, except MCF10-2A *TetOn-CENPA-FLAG-HA* cell lines stably expressing either empty vector (left) or p53-DN (right), were treated with a range of X-irradiation doses (0-15gy, X-axis). CENP-A overexpression (+ Dox (10ng/ml), green) initiated 24h prior to irradiation. Plots show mean ± 95% confidence interval for survival ratio relative to untreated p53-WT control (- Dox, black) for one experiment performed in triplicate.

C) p53-dependent X-irradiation sensitivity is specific to mid-range doxycycline levels. CFAs as in B, except using a range of doxycycline doses (0-100ng/ml, X-axis), instead of a range of X-irradiation doses. Black: unirradiated (0gy). Red: 4gy X-irradiation. Plots show mean ± 95% confidence interval for survival ratio relative to untreated p53-WT control for one experiment performed in triplicate.

D) Western blot of MCF10-2A *TetOn-CENPA-FLAG-HA* cells stably expressing either empty vector (p53-WT) or p53-DN. Total cell extracts after 24h with Dox (0, 10, or 100ng/ml, as indicated). Load: 1x, 2x, 4x, where 1x load = ~13300 cells. Primary antibodies are indicated on the right. $^{\#}$ = high sensitivity ECL exposure. H4 and γ-tubulin used as loading controls. Note that 100ng/ml Dox (referred to as 10X in Figure 5) corresponds to the Dox dose that produces the maximum CENP-A overexpression obtainable in our system, corresponding to ~2X more CENP-A protein compared to 10ng/ml Dox (referred to as 1X in Figure 5).

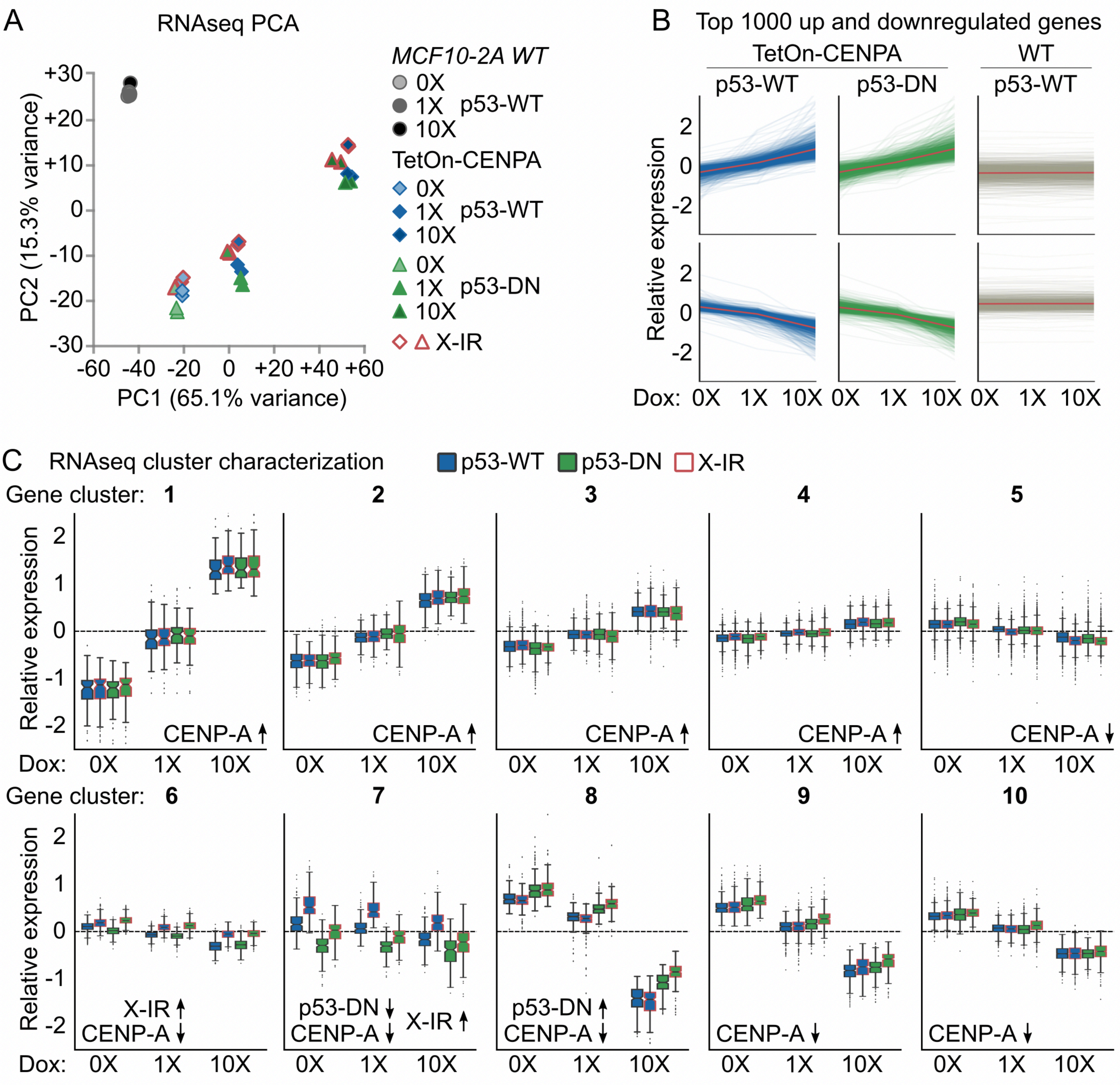
A
RNAseq PCA
MCF10-2A WT
0X
1X p53-WT
10X
TetOn-CENPA
0X
1X p53-WT
10X
0X
1X p53-DN
10X
X-IR
PC2 (15.3% variance)
PC1 (65.1% variance)
B
Top 1000 up and downregulated genes
TetOn-CENPA
WT
p53-WT
p53-DN
p53-WT
Relative expression
Dox: 0X 1X 10X 0X 1X 10X 0X 1X 10X
C
RNAseq cluster characterization
p53-WT
p53-DN
X-IR
Gene cluster: 1 2 3 4 5
Relative expression
CENP-A ↑
CENP-A ↑
CENP-A ↑
CENP-A ↑
CENP-A ↓
Dox: 0X 1X 10X
Gene cluster: 6 7 8 9 10
X-IR ↑
CENP-A ↓
p53-DN ↓
CENP-A ↓
X-IR ↑
p53-DN ↑
CENP-A ↓
CENP-A ↓
CENP-A ↓

Figure S2

**Figure S2 – Related to Figure 4 and Table S1. RNAseq non-inducible controls and gene cluster characterization**

A) Principal component analysis (PCA) of gene expression profiles from bulk RNA-seq data, as described in Figure 4A, but including non-inducible (MCF10-2A WT) controls. Each dot represents a single sample. Legend indicated below. Red outline indicates X-irradiated samples (X-IR, 4gy). PC1 on X-axis shows that most of the variance can be attributed to increasing CENP-A levels across inducible cells. Non-inducible controls cluster separately and Dox treatment does not drive major differences in their expression.

B) Effect of Dox on gene expression for inducible and non-inducible MCF10-2A cells. Relative expression from RNA-seq data of the top 1000 upregulated genes (top) and top 1000 downregulated genes (bottom) by CENP-A overexpression in MCF10-2A *TetOn-CENPA-FLAG-HA* cells, plotted by increasing Dox concentrations for each cell line. Each line represents an individual gene, while the average is shown in red for each cell line. These genes are not significantly affected by Dox in the non-inducible control.

C) Gene clusters 1-10 corresponding to heat map in Figure 4D. Box plots showing the distribution of relative expression levels of DEGs in each cluster, averaged by experimental condition. Each data point represents the mean expression level per condition, relative to average, for a particular DEG within the cluster (mean-centered counts, $\log_2$-transformed and TMM-normalized). Main effects of experimental conditions are summarized at the bottom of each plot: up arrow indicates upregulation of genes in the cluster with CENP-A overexpression, X-irradiation, or p53-DN; down arrow indicates downregulation. Clusters 1 to 4 comprise genes that are upregulated as CENP-A is overexpressed, while genes in Clusters 5 to 10 are downregulated, each to varying degrees. Interestingly, genes in Clusters 6 to 8 also showed a coordinated response to radiation treatment or p53 inactivation. Importantly, the only cluster where p53 inactivation showed opposite effects to CENP-A overexpression was Cluster 8. Cluster 8 is also displayed in Figure 4E, duplicated here for direct comparison with other clusters. The top 10 enriched KEGG pathways for each DEG cluster are provided in Table S1.

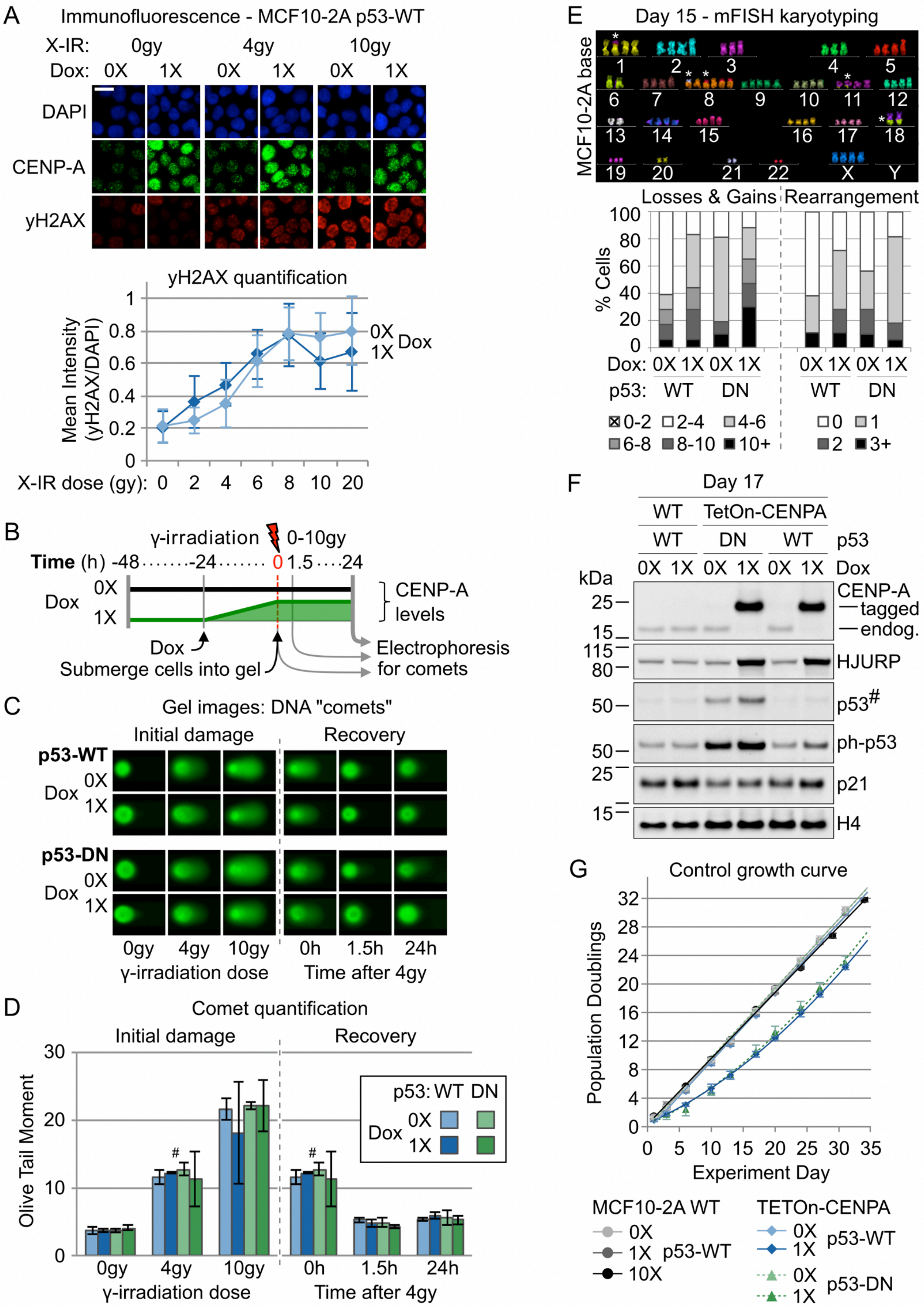
A
Immunofluorescence - MCF10-2A p53-WT
X-IR: 0gy 4gy 10gy
Dox: 0X 1X 0X 1X 0X 1X
DAPI
CENP-A
yH2AX
yH2AX quantification
Mean Intensity (yH2AX/DAPI)
0X 1X Dox
X-IR dose (gy): 0 2 4 6 8 10 20
B
γ-irradiation 0-10gy
Time (h) -48 -24 0 1.5 24
Dox 0X 1X
CENP-A levels
Dox
Submerge cells into gel
Electrophoresis for comets
C
Gel images: DNA "comets"
Initial damage
Recovery
p53-WT
p53-DN
Dox 0X 1X
0gy 4gy 10gy
0h 1.5h 24h
γ-irradiation dose
Time after 4gy
D
Comet quantification
Initial damage
Recovery
Olive Tail Moment
p53: WT DN
Dox 0X 1X
0gy 4gy 10gy
0h 1.5h 24h
γ-irradiation dose
Time after 4gy
E
Day 15 - mFISH karyotyping
MCF10-2A base
Losses & Gains
Rearrangement
% Cells
Dox: 0X 1X 0X 1X 0X 1X 0X 1X
p53: WT DN WT DN
0-2 2-4 4-6 6-8 8-10 10+
0 1 2 3+
F
Day 17
WT TetOn-CENPA
WT DN WT p53
0X 1X 0X 1X 0X 1X Dox
kDa
CENP-A
tagged
endog.
HJURP
p53#
ph-p53
p21
H4
G
Control growth curve
Population Doublings
Experiment Day
MCF10-2A WT
0X
1X p53-WT
10X
TETOn-CENPA
0X 1X p53-WT
0X 1X p53-DN

Figure S3

**Figure S3 – Related to Figures 4 and S1. Impact of CENP-A overexpression of cell proliferation, DNA damage and CIN**

A) γH2AX response of p53-WT cells after X-irradiation with or without CENP-A overexpression. MCF10-2A *TetOn-CENPA-FLAG-HA* cells with empty vector (p53-WT) were plated in culture dishes containing collagen/fibronectin-coated glass cover slips and then treated with 10ng/ml Dox or no Dox control for 24h. Cells were then irradiated by X-ray generator with 0-20gy and immediately fixed for immunofluorescence with γH2AX and CENP-A antibodies. Top: representative max intensity projection images from Z-stack for cells with (+) or without (-) 10ng/ml Dox after 0, 4 or 10gy X-irradiation. Bottom: quantification of mean γH2AX/DAPI intensity per nucleus for each condition (ImageJ) ± Standard Deviation. N = >90 for each condition.

B) Scheme of comet assays with relative CENP-A protein levels over time for MCF10-2A *TetOn-CENPA-FLAG-HA* cell lines. To test the impact of p53 status and CENP-A overexpression on DNA damage directly, we performed alkaline comet assays after γ-irradiation on cells stably expressing empty vector (p53-WT) or p53-DN, with (+) or without (-) 24h 10ng/ml Dox.

C) Representative images of comets pertaining to scheme in B.

D) Quantification of DNA damage measured as the product of the comet tail length and fraction of DNA in tail (olive tail moment) for each condition pertaining to scheme in B and representative images in C. Plots show mean ± standard deviation for the mean olive tail moment from three independent experiments. [#]Same data displayed for 4gy 0h treatment shown in both plots for comparison.

E) Quantification of chromosomal abnormalities from mFISH karyotypes in MCF10-2A *TetOn-CENPA-FLAG-HA* cells with either empty vector (p53-WT) or dominant-negative p53 (p53-DN) grown continuously for 15 days without Dox (0X) or with Dox (1X, 10ng/ml). Top: Representative Metafer automated image of chromosome spread from MCF10-2A *TetOn-CENPA-FLAG-HA* cells, empty vector (p53-WT) without Dox. Asterisks denote baseline chromosome rearrangements that were present in all p53-WT no Dox spreads. Bottom: Stack bar plots showing percentage of binned counts from at least 17 metaphase spreads per condition pertaining to legends below. Left: Losses and gains of chromosomes per cell determined as the sum of the difference from the mean for each chromosome of the p53-WT no Dox control. Right: New chromosome rearrangements per cell were determined as the total number of structural chromosomal anomalies observed per spread, excluding the ones that were common to all p53-WT no Dox spreads.

F) Western blot of TCEs corresponding to Day 17 of growth curve in G for 0X and 1X Dox conditions. Load corresponds to ~50000 cells each. Primary antibodies are indicated on the right. [#] = high sensitivity ECL exposure. H4 used as loading control.

G) Growth curve, as in Figure 4G, but including the non-inducible parental (MCF10-2A WT) cells. Growth curves of MCF10-2A *TetOn-CENPA-FLAG-HA* cells from 0X and 1X Dox (Figure 4G) shown for comparison. Legend: Non-inducible, grey circles; p53-WT, blue diamonds; p53-DN, green triangles. Increasing darkness of fill corresponds to increasing Dox 0X, 1X (10ng/ml) or 10X (100ng/ml).

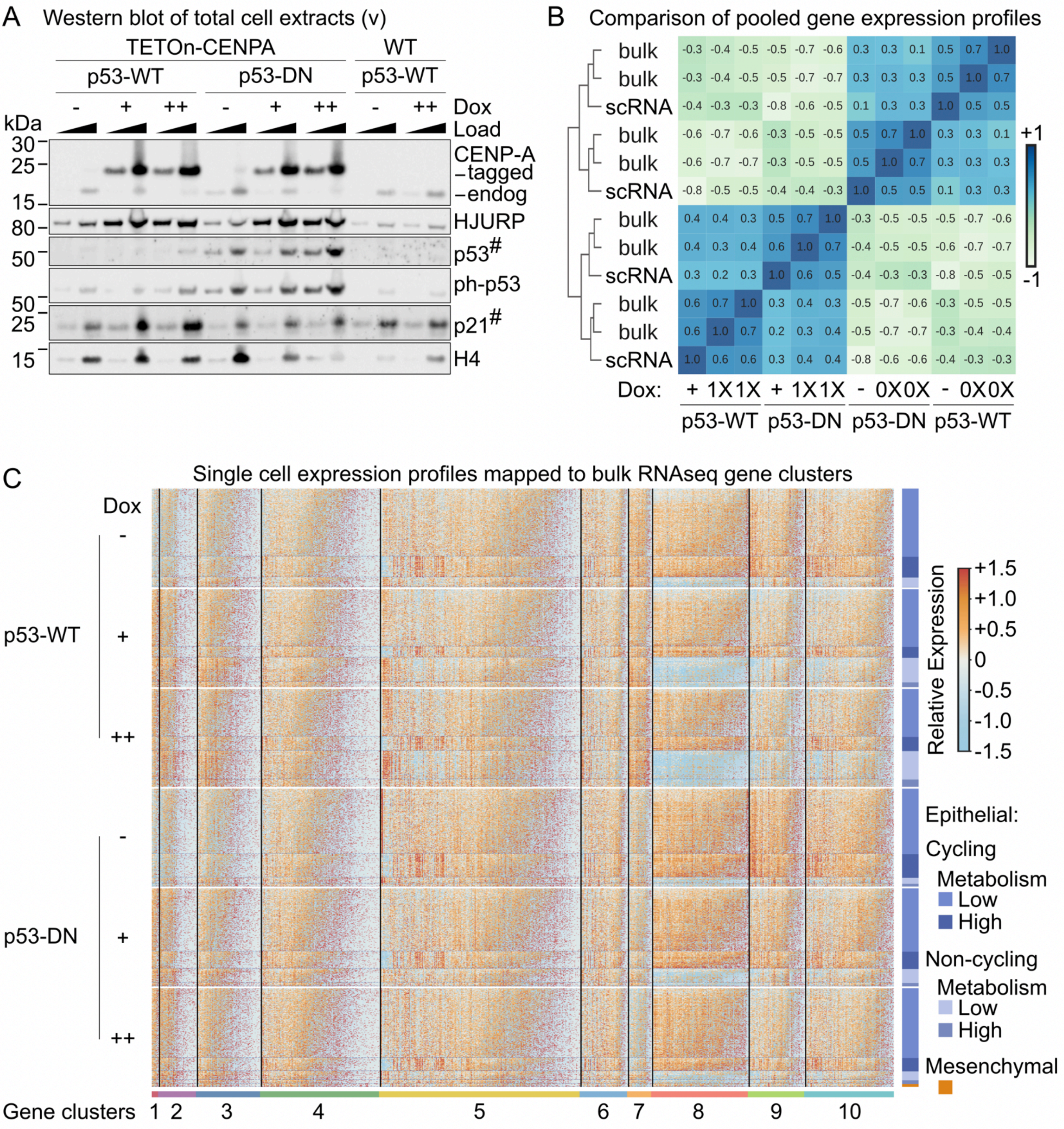
A
Western blot of total cell extracts (v)
TETOn-CENPA
WT
p53-WT
p53-DN
p53-WT
- + ++ - + ++ - ++ Dox
Load
kDa
30
25
15
80
50
50
25
15
CENP-A
–tagged
–endog
HJURP
p53#
ph-p53
p21#
H4
B
Comparison of pooled gene expression profiles
bulk -0.3 -0.4 -0.5 -0.5 -0.7 -0.6 0.3 0.3 0.1 0.5 0.7 1.0
bulk -0.3 -0.4 -0.5 -0.5 -0.7 -0.7 0.3 0.3 0.3 0.5 1.0 0.7
scRNA -0.4 -0.3 -0.3 -0.8 -0.6 -0.5 0.1 0.3 0.3 1.0 0.5 0.5
bulk -0.6 -0.7 -0.6 -0.3 -0.5 -0.5 0.5 0.7 1.0 0.3 0.3 0.1
bulk -0.6 -0.7 -0.7 -0.3 -0.5 -0.5 0.5 1.0 0.7 0.3 0.3 0.3
scRNA -0.8 -0.5 -0.5 -0.4 -0.4 -0.3 1.0 0.5 0.5 0.1 0.3 0.3
bulk 0.4 0.4 0.3 0.5 0.7 1.0 -0.3 -0.5 -0.5 -0.5 -0.7 -0.6
bulk 0.4 0.3 0.4 0.6 1.0 0.7 -0.4 -0.5 -0.5 -0.6 -0.7 -0.7
scRNA 0.3 0.2 0.3 1.0 0.6 0.5 -0.4 -0.3 -0.3 -0.8 -0.5 -0.5
bulk 0.6 0.7 1.0 0.3 0.4 0.3 -0.5 -0.7 -0.6 -0.3 -0.5 -0.5
bulk 0.6 1.0 0.7 0.2 0.3 0.4 -0.5 -0.7 -0.7 -0.3 -0.4 -0.4
scRNA 1.0 0.6 0.6 0.3 0.4 0.4 -0.8 -0.6 -0.6 -0.4 -0.3 -0.3
+1
-1
Dox: + 1X1X + 1X1X - 0X0X - 0X0X
p53-WT p53-DN p53-DN p53-WT
C
Single cell expression profiles mapped to bulk RNAseq gene clusters
Dox
p53-WT
-
+
++
p53-DN
-
+
++
Relative Expression
+1.5
+1.0
+0.5
0
-0.5
-1.0
-1.5
Epithelial:
Cycling
Metabolism
Low
High
Non-cycling
Metabolism
Low
High
Mesenchymal
Gene clusters 1 2 3 4 5 6 7 8 9 10

Figure S4

**Figure S4 – Related to Figures 4, 5, 6 and S5. Single-cell RNA-seq support and comparison to bulk RNA-seq**

A) Western blot of total cell extracts pertaining to time point 'v' in Figure 5A, including MCF10-2A non-inducible (WT) parental control cells after long-term exposure to Dox. All cells passaged in parallel for 69 days, or 55 days in case of parental line. Number of days exposed to 1X Dox (10ng/ml) indicated above. Aliquots from the same samples were used for the scRNA-seq experiments. Primary antibodies are indicated on the right. Load: 1x, 3x; where 1x = ~16700 cells. $^{\#}$ = high sensitivity ECL exposure. H4 used as loading control.

B) Heat map: clustering of pooled scRNA-seq and bulk RNA-seq samples from comparable experimental conditions based on Pearson correlation among gene expression profiles. Left: row dendrogram showing the clustering of the samples based on average linkage. Corresponding p53 status and Dox treatment indicated on the X-axis (+ or 1X = 10ng/ml; - or 0X = no Dox; for scRNA-seq and bulk RNA-seq respectively). Positive correlations (blue) and negative correlations (white) are indicated by intensity.

C) Heat map of single-cell expression profiles according to bulk RNA-seq DEG clusters (1-10, labeled at the bottom). 300 cells were randomly sampled per experimental condition. Each row represents a single cell, ordered by experimental condition (indicated on the left) and scRNA-seq cell cluster (indicated on the right, see Figure S5A and Table S2 for cell cluster identification). Within each cluster, cells are sorted based on total counts. Each column represents a DEG identified in the bulk RNA-Seq results, ordered by gene cluster and sorted by average expression across all cells. The colour gradient is proportional to the relative expression level across individual cells, from low (blue) to high (orange).

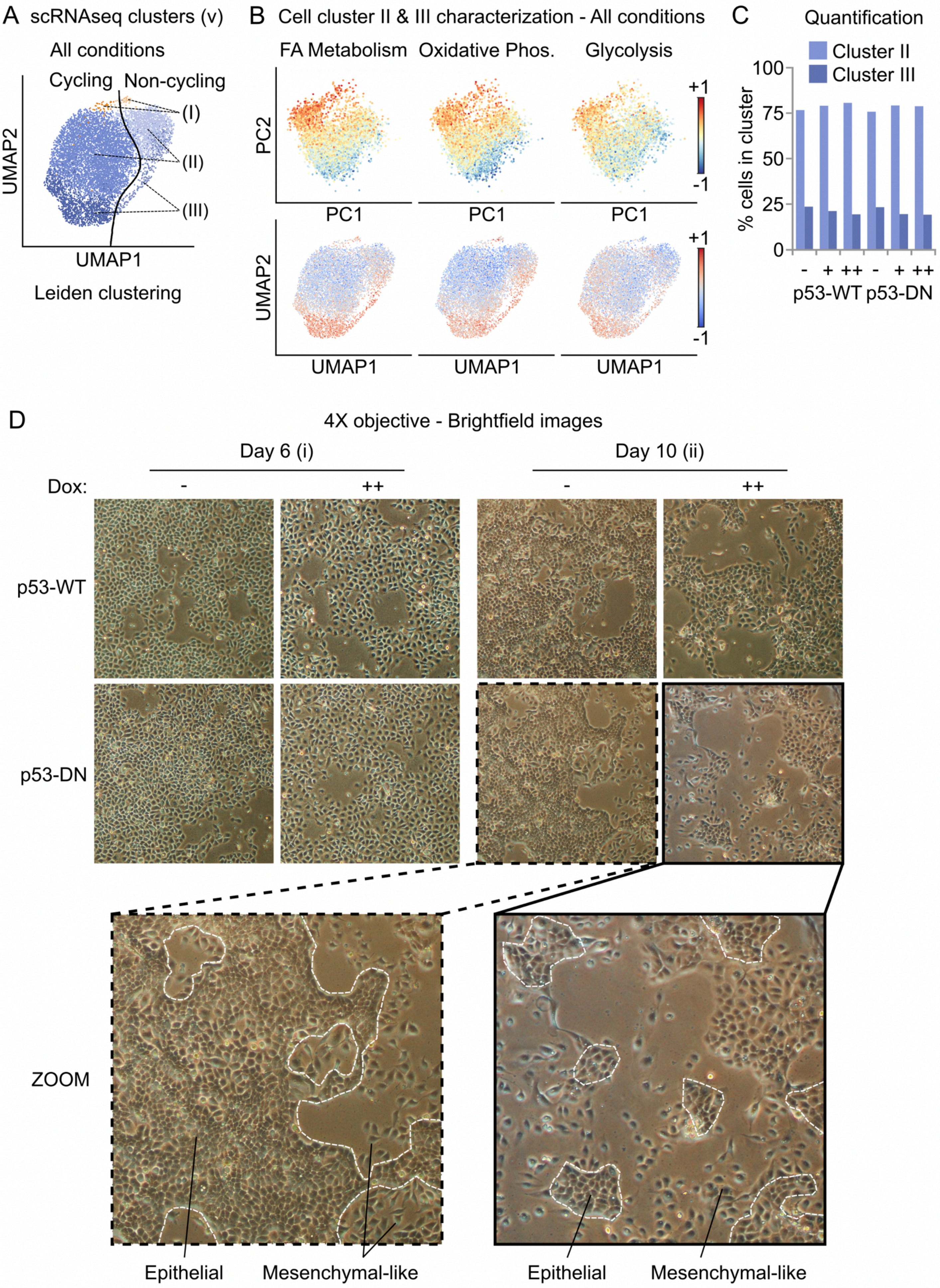
A scRNAseq clusters (v)
All conditions
Cycling Non-cycling
(I)
(II)
(III)
UMAP2
UMAP1
Leiden clustering
B Cell cluster II & III characterization - All conditions
FA Metabolism Oxidative Phos. Glycolysis
PC2
PC1
+1
-1
UMAP2
UMAP1
C Quantification
Cluster II
Cluster III
% cells in cluster
100
75
50
25
0
- + ++ - + ++
p53-WT p53-DN
D 4X objective - Brightfield images
Day 6 (i)
Day 10 (ii)
Dox:
-
++
p53-WT
p53-DN
ZOOM
Epithelial
Mesenchymal-like

Figure S5

**Figure S5 – Related to Figures 5, 6 and S4 and Table S2. Single-cell RNA-seq epithelial cell clusters and emergence of mesenchymal-like characteristics by brightfield microscopy**

A) UMAP plots of scRNA-seq data from time point ‘v’ in Figure 5A, showing three main clusters of cells based on Leiden clustering after cell cycle correction. Cluster I (orange), Cluster II (light blue), and Cluster III (dark blue), are sub-divided as either Cycling (dark) or Non-cycling (pale), according to the previous analysis in Figure 5. See Table S2 for gene markers associated with each cluster, ranked by one-vs-all logistic regression. Cluster I is negative for epithelial markers (e.g. *CDH1*, *EPCAM* and *KRT19*) and shows high expression of mesenchymal genes (e.g. *VIM*, *FN1* and *CDH2*). Clusters II and III are positive for epithelial markers but show differential expression of genes involved in cell metabolism (e.g. *CTSV, AFMID, GLA, ACAT2, CBR4*).

B) Top: Cell-to-cell variability along the second principal component of the scRNA-seq data can be explained by differences in expression of cell metabolism genes. Principal component analysis (PCA) of scRNA-seq experiments at time point ‘v’ in Figure 5A. All conditions merged. Each dot represents a single cell on the first (PC1) and second (PC2) principal component, coloured by expression of genes involved in Fatty Acid (FA) Metabolism, Oxidative Phosphorylation (Phos.), and Glycolysis (Hallmark gene sets, MSigDB v6.2). Bottom: As above, except data plotted in UMAP space. Epithelial clusters II and III show low and high expression of genes involved in cell metabolism processes, respectively.

C) Bar plots showing percentage of scRNA-seq Clusters II (cell metabolism low) and III (cell metabolism high) in A by experimental condition. Proportions are similar for untreated p53-WT and p53-DN cells. CENP-A overexpression (acute or chronic) reduces the proportion of cell metabolism high cells in both cases and the opposite for cell metabolism low cells. We speculate that this represents natural cell-to-cell variation in cell metabolism associated with nutrient availability in the cells, which may be hindered by increased production of CENP-A.

D) Brightfield images of cells at time points ‘i’ and ‘ii’ in Figure 5A. White dashed lines in zoomed images outline edges of epithelial cell groups. Epithelial cells are characterized by strong cell-cell contacts, compared to cells with mesenchymal-like characteristics (isolated/showing reduced cell-cell contacts and elongated/distorted shape). Images taken of live cells by brightfield microscopy with a 4X objective.

Supplementary Tables

**Table S1 - Related to Figures 4 and S2. Bulk RNA-seq data analysis of MCF10-2A cells after CENP-A overexpression, switch of p53 status, and X-irradiation**

Sheet 1: Contents & design table
Sheet 2: Normalized counts (excluding non-inducible parental control (WT))
Sheet 3: Normalized counts (including WT)
Sheet 4: Differential expression results of all genes according to CENP-A overexpression (OE via Dox) effect, p53 effect (p53-DN), and/or X-irradiation
Sheet 5: All genes ranked according to CENP-A overexpression effect (OE via Dox) based on fold change and p-value
Sheet 6: Gene Set Enrichment Analysis by overall CENP-A overexpression effect (OE via Dox) - KEGG pathways, WebGestaltR v0.4.2
Sheet 7: Over-Representation Analysis per gene cluster - KEGG pathways, WebGestaltR v0.4.2

**Table S2 – Related to Figures 6, S4 and S5. Single-cell RNA-seq cell cluster gene markers**

Sheet 1: Contents
Sheet 2: Mesenchymal/EMT high (all) vs rest
Sheet 3: Epithelial Cell Metabolism Low (all) vs rest
Sheet 3: Epithelial Cell Metabolism High (all) vs rest
Sheet 4: Mesenchymal/EMT high (cycling) vs rest
Sheet 5: Mesenchymal/EMT high (non-cycling) vs rest
Sheet 6: Epithelial Cell Metabolism Low (cycling) vs rest
Sheet 7: Epithelial Cell Metabolism Low (non-cycling) vs rest
Sheet 8: Epithelial Cell Metabolism High (cycling) vs rest
Sheet 9: Epithelial Cell Metabolism High (non-cycling) vs rest
